# Nuclear-injuries by aberrant dynein-forces defeat proteostatic purposes of Lewy Body Inclusions

**DOI:** 10.1101/2021.10.29.466393

**Authors:** Shemin Mansuri, Aanchal Jain, Richa Singh, Shivali Rawat, Debodyuti Mondal, Swasti Raychaudhuri

## Abstract

Biogenesis of inclusion bodies (IBs) facilitates protein quality control (PQC). Canonical aggresomes execute degradation of misfolded proteins while non-degradable amyloids quarantine into Insoluble Protein Deposits. Lewy Bodies (LBs) are well-known neurodegenerative IBs of α-Synuclein associated diseases like Dementia with Lewy bodies (DLB) and Parkinson’s disease. Intriguingly, the PQC benefits and drawbacks associated with LBs remain underexplored. Here, we report that the crosstalk between LBs and aggresome-like IBs of α-Synuclein (Syn-aggresomes) buffers amyloidogenic α-Synuclein load in primary neurons and mitotic cell models. Using cellular biochemistry, genetic knockdown, and microscopy tools we find that LBs possess unorthodox PQC-capacities of self-quarantining the Syn-amyloids, and being degradable upon receding fresh amyloidogenesis. Syn-aggresomes equilibrate biogenesis of LBs by facilitating spontaneous and opportunistic turnover of soluble α-Synuclein and Syn-amyloids, respectively. However, LBs overgrow at the perinucleus once amyloidogenesis sets in and are misidentified by cytosolic BICD2 as cargos for motor-protein dynein. Simultaneously, microtubules surrounding the perinuclear LBs are distorted, misbalancing the dynein motor-force on nucleoskeleton leading to lamina injuries. Like typical Laminopathies, nucleocytoplasmic mixing, DNA-damage, and deregulated transcription of stress chaperones defeat the proteostatic purposes of LBs. We confirmed the lamina disintegrities in brain sections of Parkinson’s disease patients. Together, our study provides insights into the intricate complexities of proteostatic possibilities associated with α-Synuclein inclusions and offers understanding on the proteostasis-sensitivity of LB-containing aging neurons via lamina-injuries.

**Significance statement:** Amyloid inclusions of α-Synuclein called Lewy Bodies (LBs) are hallmark of multiple neurodegenerative diseases like Dementia with Lewy Bodies and Parkinson’s disease. A significant reason for the degeneration of LB-containing aging neurons in these diseases is their sensitivity to proteostasis stresses. We find two distinct inclusions of α-Synuclein in the same neurons/cells. First, the Syn-aggresomes titrate the biogenesis of the other i.e., the LBs, by facilitating degradation of soluble α-Synuclein. Syn-amyloids deposited in LBs remain degradable but LBs overgrow at the perinucleus once the kinetics of amyloid-biogenesis exceeds aggresome-assisted degradation of α-Synuclein. Perinuclear LBs destabilize local tubulin-meshwork and are misidentified as cargo for cytosolic motor dynein. Aberrant cytoskeleton-nucleoskeleton tension disintegrates lamina, deregulates stress-responsive transcription, and triggers proteostasis-sensitivity in LB-containing neurons.

## Introduction

Amyloid inclusion bodies (IBs) are hallmark of several age-related neurodegenerative disorders. The highly organized and compact IBs of amyloids are commonly known as Insoluble Protein Deposits or IPODs. Many amyloidogenic proteins accumulate into IPODs due to their intrinsic property of self-assembly and inefficient ubiquitination (1–3). Amyloid-IBs can be degenerative (4, 5). Contradictory evidences suggest that rapid IPOD-biogenesis is intended to protect cellular homeostasis by limiting promiscuous degenerative interactions of the oligomeric amyloids with other proteins (2, 6). IPODs may be peripheral or perinuclear depending on cytoskeleton association, can be inefficiently degraded by autophagy, and are asymmetrically inherited between the daughter cells during mitosis (1, 7, 8). Occasionally, amyloidogenic proteins also accumulate into perinuclear aggresome or juxtanuclear quality control compartment (JUNQ) which serve as localized sites for ubiquitin-proteasome system (UPS) mediated degradation (9, 10). This delays the UPS-mediated turnover of other misfolding-prone proteins in the same cells (10).

Strikingly, homeostatic characterization of the amyloid IBs is so far limited to only few candidate amyloid proteins like mutant Huntingtin (mHtt) and prion proteins Rnq1, Ure2, Sup35 etc., which spontaneously form compact IPOD-like IBs in mitotic models. In contrast, the Lewy Bodies (LBs), the perinuclear loosely-packed filamentous IBs of amyloidogenic α-Synuclein observed in Parkinson’s disease (PD) (11), are inconvenient to model and remained underexplored. The canonical aggresomes are positioned at the microtubule organizing centre (MTOC) (12). LBs stained by ubiquitin and other PQC-factors in PD post-mortem brains are also marked by MTOC marker γ-tubulin and therefore claimed as ‘failed aggresomes’ (13, 14). While MTOCs are visible as dots of γ-tubulin at the perinucleus (15), the size of the γ-tubulin-positive IBs in PD brains are even bigger than the cell nucleus (13). Further, UPS-mediated protein turnover potential at the LBs has never been verified.

Here, we report that LBs are not aggresomes but extra-aggresomal filamentous IBs of α-Synuclein. Canonical aggresomes of α-Synuclein (Syn-aggresomes) coexist in the same cells which represent a spontaneous homeostatic response of cellular-PQC to sequester and degrade α-Synuclein. In contrast, LBs offer an unorthodox self-quarantine-strategy for α-Synuclein amyloids to facilitate their opportunistic degradation rather than terminally aggregating the amyloids into conventional, poorly degradable IPODs. Paradoxically, impaired transcription of stress-chaperones due to nuclear lamina injuries is a late degenerative outcome in cells with overgrown perinuclear LBs that imposes cellular vulnerability upon proteostasis stresses. We identify that a dynein-based misbalanced cytoskeleton force working on the nucleoskeleton is responsible for the lamina disintegration. Remarkably, engineered conversion of perinuclear LB-like IBs into intranuclear Syn-filaments protects lamina-integrity as well as stress-transcription suggesting lamina-injuries as a major driver of proteostasis sensitivity in LB-containing cells. Lamina injuries are widespread in post-mortem brain sections of PD patients suggesting α-Synuclein proteinopathies may also be considered as Laminopathies.

## Results

### Two unique inclusion bodies of α-Synuclein in primary neurons

α-Synuclein deposits into Lewy Body (LB) like filamentous inclusions in primary neurons upon seeding with preformed amyloid fibrils (16, 17). These IBs are stained by phospho-Ser129-Synuclein (p129-Syn) antibody. Compared to ∼90% of phosphorylation in LBs, only ∼4% of α-Synuclein is phosphorylated in healthy neurons and thereby very poorly or not visible by microscopy (18–20). To detect if phosphorylated α-Synuclein was deposited into any inclusion bodies (IBs) in unseeded neurons, we stained the primary neurons isolated from hippocampus of E19 pre-natal C57BL/6 mice using phospho-Ser129-Synuclein (p129-Syn) antibody ((MJF-R13 (8–8), epitope: phosphoserine 129) and imaged at higher gain of laser power in a confocal microscope. We observed continuous staining of p129-Syn at the neuronal projections and light dotted bodies at the periphery and inside nucleus (**Fig. 1Ai**, **MIP**). γ-Tubulin staining at the centrosomes was observed just outside the nucleus (**Fig. S1A**). We also observed blobs of p129-Syn positioned on the perinuclear γ-tubulin staining (**Fig. 1Ai and S1Bi**, stick arrowheads). Intriguingly, staining the neurons with another p129-Syn antibody, P-syn/81A, epitope: phosphoserine 129, resulted in only dotted appearance of p129-Syn at the cell body (**Fig. 1Bi**). Perinuclear blobs of p129-Syn co-stained with autophagosome cargo marker Sequestosome 1 (SQSTM-1/p62) were also observed in these neurons (**Fig. 1Bi and S1Ci**, stick arrowheads). Notably, both γ-tubulin and SQSTM-1 are known markers of canonical aggresomes (21).

**Fig 1.**
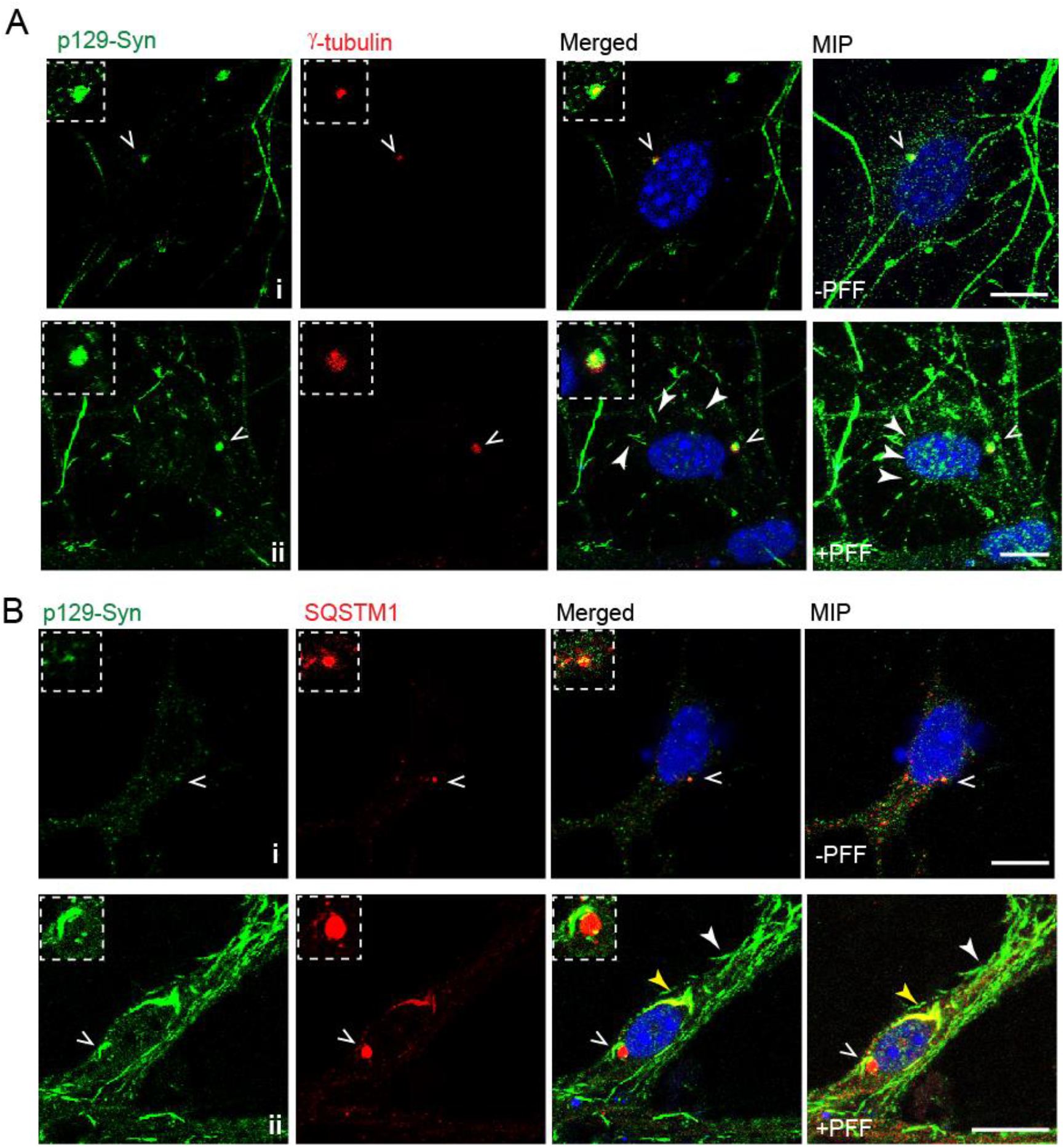
Two unique inclusion bodies of α-Synuclein in primary neurons. **A**: Primary neurons, with and without 100 nM PFF treatment from 5 days *in vitro* (DIV) and immunostained at 15 DIV with p129-Syn (MJF-R13 (8-8), epitope: phosphoserine 129) and γ-tubulin antibodies. **B:** Primary neurons with and without PFF treatment from 5 DIV immunostained at 15 DIV with p129-Syn (P-syn/81A, epitope: phosphoserine 129) and SQSTM1 antibodies. Arrowheads indicating different kind of inclusions. Insets: zoomed images of the indicated locations. Scale bar-10 µm. Different channels of single z-plane and Maximum Intensity Projection (MIP) of the same field is shown.

Next, we treated the hippocampal neurons with recombinant α-Synuclein fibrils (preformed fibrils, PFF). PFF treatment for 10 days resulted in accumulation of α-Synuclein in filamentous structures (Syn-filaments) (**Fig. 1Aii–Bii and S1Bii–Cii**). The continuous staining of p129-Syn at the projections of unseeded neurons (**Fig. 1Ai and SBi**) was converted into a fragmented appearance (**Fig. 1Aii and SBii**). We observed several neurons with smaller Syn-filaments at the projections and close to the nucleus (**Fig. 1Aii**, white arrowheads **and S1Bii**). Deposition of multiple filaments at the perinucleus resulted in biogenesis of larger LB-like IBs (**Fig. 1Bii**, yellow arrowhead) morphologically similar to pathological LBs (19). Smaller filaments were also present at the peripheral cell body and projections of the same neurons (**Fig. 1Bii**, white arrowhead). The LB-like perinuclear Syn-IBs were partially co-stained by SQSTM-1 while small Syn-filaments at the peripheral remained unrecognized by SQSTM-1 antibody (**Fig. 1Bii**). Remarkably, p129-Syn blobs positioned on the perinuclear γ-tubulin were visible in the primary neurons with smaller Syn-filaments (**Fig. 1Aii**). Similar γ-tubulin positive p129-Syn blobs was difficult to identify in large perinuclear LB-like IB containing neurons but we did observe SQSTM-1 blobs surrounded by p129-Syn staining independent of the large LB-like IBs (**Fig. 1Bii and S1Cii**).

LBs engulf and distort membranous organelles like endoplasmic reticulum (ER) and mitochondria (19, 22). Calnexin and CMX-Ros staining revealed distribution of ER and mitochondria throughout the cell body and the projections of the unseeded primary neurons (**Fig. S1Di and S1Ei**). Like the LBs in patients, the LB-like perinuclear IBs in our experiments partially enwrapped both ER and mitochondria and their smooth distribution throughout the neurons was lost (**Fig. S1Dii–Eii**). Thus, we captured two distinct IBs in the same neurons mimicking the perinuclear aggresomes and perinuclear LBs into which α-Synuclein was deposited simultaneously.

### Co-existence of two α-Synuclein IBs is consistent in mitotic cells

While LB-like filamentous IBs of α-Synuclein could be modelled in post-mitotic neurons, similar inclusions are not reported in mitotic cells. Nevertheless, aggresome-like Syn-IBs seem to appear by manipulating culture conditions in α-Synuclein overexpressing cells (23–28). Western blots indicated undetectable load of α-Synuclein in Hek293T cells (**Fig. 2A**). Therefore, we exogenously expressed α-Synuclein in Hek293T cells to address two questions. First, can amyloidogenic α-Synuclein form LB-like filamentous IBs in mitotic cells? Second, can the non-native α-Synuclein be partitioned into two distinct IBs in Hek293T cells likewise the endogenous α-Synuclein in primary neurons?

**Fig. 2:**
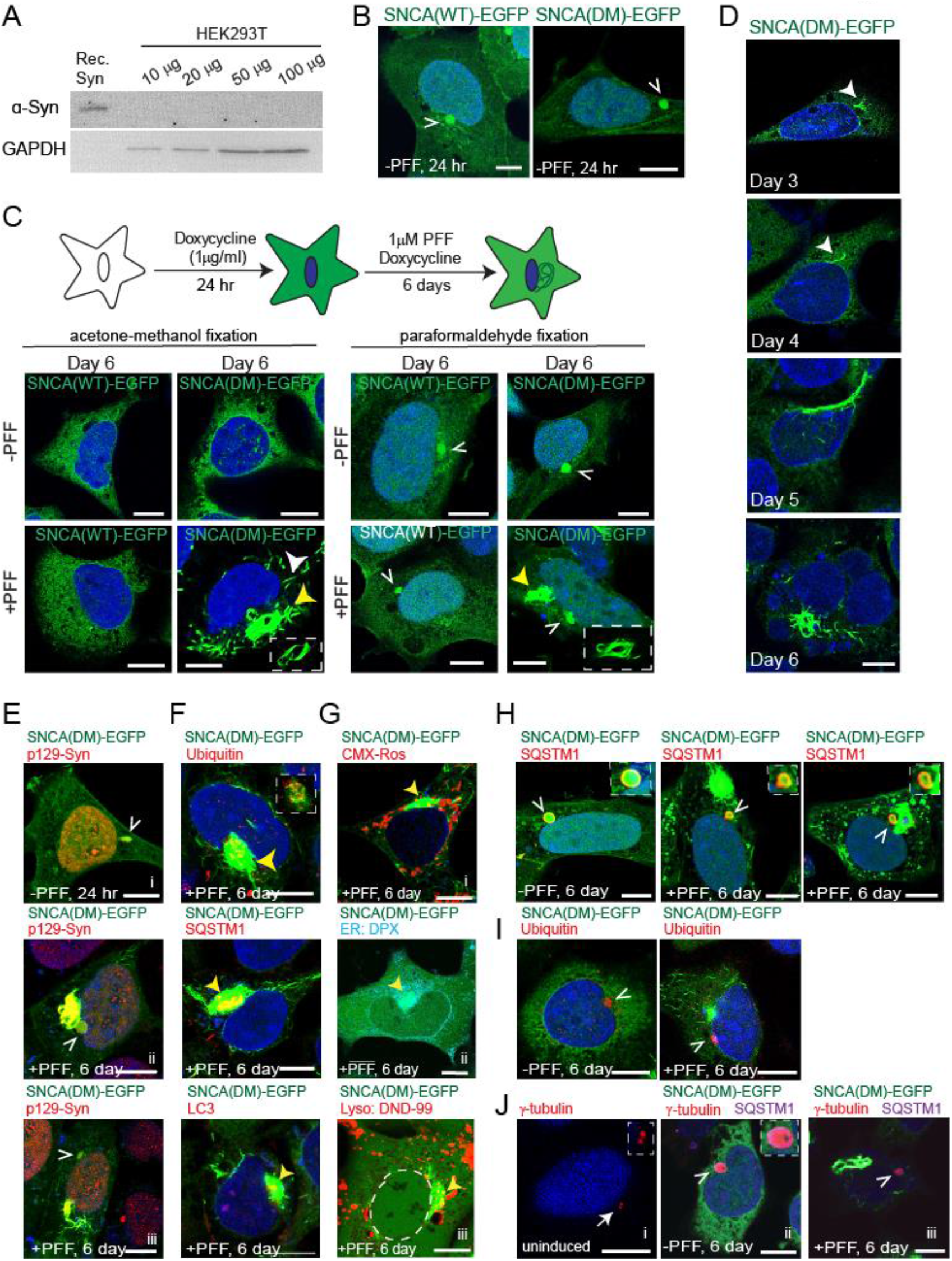
Co-existence of two α-Synuclein IBs is not restricted to post-mitotic neurons. **A:** Increasing concentration of Hek293T cell lysate immunoblotted for total α-Synuclein (α-Syn) (14H2L1; epitope:117-125). Lane 1-recombinant α-Synuclein control. GAPDH as loading control. **B:** Hek293T stable lines expressing EGFP-tagged WT and DM (A30P+A53T) α-Synuclein (SNCA). Cells were imaged after 24 hr of expression. Stick arrowheads indicate SNCA-EGFP blob. **C: Top:** Experimental scheme. **Bottom**: Hek293T stable lines expressing EGFP-tagged α-Synuclein (SNCA) WT and DM. Cells were fixed with Acetone-Methanol (left panel) or PFA (right panel) after 6 days with or without PFF treatment and imaged. Stick arrowheads indicate SNCA-EGFP blob. Solid white and yellow arrowheads indicate small filaments and LB-like filaments respectively. Insets: low exposure images of the yellow arrowhead indicated regions. **D:** Localization of Syn-filaments in SNCA(DM)-EGFP cells at different days of PFF-incubation. Solid white arrowheads indicate small filaments. **E:** 6^th^ day PFF-treated and untreated SNCA(DM)-EGFP cells were immunostained with p129-Syn (MJF-R13 (8-8)) antibody and imaged. Stick arrowheads in **E i, ii and iii** indicate a separate blob stained by p129-Syn antibody. **F:** 6^th^ day PFF-treated SNCA(DM)-EGFP cells were immunostained with markers for canonical LBs and imaged. Yellow arrowheads indicate co-localization. Inset – low exposure image of arrow indicated region in Ubiquitin antibody-stained cell. **G:** 6^th^ day PFF-treated SNCA(DM)-EGFP cells were stained for various cell organelles and imaged. Dashed circles indicate nuclear boundary in **Giii**. Yellow arrowheads indicate co-localization. **H:** 6^th^ day PFF-treated and untreated SNCA(DM)-EGFP cells immunostained with SQSTM1 antibody. Insets – zoomed images of the arrow indicated locations. **I:** 6^th^ day PFF-treated and untreated SNCA(DM)-EGFP cells immunostained with ubiquitin antibody. Stick arrowhead indicates the perinuclear blob. **J:** Uninduced and dox-induced 6^th^ day SNCA(DM)-EGFP cells were immunostained with SQSTM1 and γ-tubulin antibodies. Stick arrowheads indicate SNCA-EGFP blob. Insets: zoomed images of the arrow indicated regions. Scale bar – 10 µm. **Fig. 2B, E, H** – Paraformaldehyde fixation. **Fig. 2D, F, Gi, I, J** – Acetone:Methanol fixation. **Fig. 2Gii–iii** – live cell microscopy without fixation.

We prepared stable inducible lines in Hek293T cells exogenously expressing α-Synuclein with EGFP at the C-terminus (SNCA-EGFP). Wild type SNCA-EGFP was diffusely distributed throughout the cell including the nucleus. A perinuclear SNCA-EGFP blob was also detected within 24 hr of induction (**Fig. 2B**; stick arrowhead). Filamentous IBs of wild type SNCA-EGFP was not triggered even upon prolonged seeding with 1 μM PFF (**Fig. 2C**). Two mutations, A30P and A53T, in α-Synuclein are associated with familial PD (29, 30). Intriguingly, a double mutant A30P+A53T (SNCA(DM)) recapitulates aggressive early onset degenerative phenotypes of Synucleopathies in mice models (31, 32).

Similar to wild type, A30P, A53T, and DM-EGFP variants were present throughout the Hek293T cells (**Fig. 2C and S2A**). Among these, A53T and DM-EGFP expressing cells showed Syn-filaments after prolonged seeding (**Fig. 2C and S2A**, yellow arrowheads). Although non-significant, number of Syn-filament containing cells was more in case of SNCA(DM)-EGFP than SNCA(A53T)-EGFP (**Fig. S2B**). Perinuclear blobs of SNCA(DM)-EGFP were also present irrespective of the occurrence of Syn-filaments (**Fig. 2C**, right panel, stick arrowheads). While these blobs were prominent in paraformaldehyde-fixed cells, Syn-filament morphologies were better visible after acetone-methanol fixation. This indicated differences in fluidity and cross-linking probability of α-Synuclein molecules deposited in these two co-existing IBs (33). We generated a SNCA(DM) stable line in Hek293T without the EGFP tag which also formed LB-like inclusions ruling out any effect of C-terminal EGFP-tagging on Syn-filament biogenesis (**Fig. S2C**).

We observed multiple initiation sites for Syn-filament biogenesis (**Fig. 2D**, **S2D and Supplementary Video 1**). Small Syn-filaments first appeared in the cytoplasm after 3 days of seeding followed by formation of bundle of filaments. These bundle of filaments gradually merged into skein-like filamentous IBs encaging the nucleus. The 6^th^ day cell population was heterogeneous showcasing different stages of filament biogenesis. Large skein-like IBs were mostly perinuclear and were observed in more than 60 % of the Syn-filament-containing cells (**Fig. S2E**). Syn-filaments were also infrequently observed inside or distant to the nucleus (**Fig. S2Fii–iii**). Thus, the progression of Syn-filament biogenesis in the cell culture model was identical with primary neurons where small filaments are first observed at the projections and peripheral cell body that gradually converge into perinuclear LB-like IBs (19). Syn-filament biogenesis was prevented by Congo Red (CR) without affecting cell viability **(Fig. S2G–H**), a beta sheet binding dye known to prevent *in vitro* fibrilization of α-Synuclein by inhibiting oligomerization (34–36).

Like primary neurons, smaller or larger perinuclear Syn-IBs were always positive for p129-Syn antibody in our Hek293T cell culture model (**Fig. 2Eii–iii and S3A**). Ubiquitin, autophagosome cargo marker Sequestosome 1 (SQSTM-1/p62) and LC3 antibodies stained only the large perinuclear filamentous IBs at the core but the smaller peripheral filaments remained unrecognized (**Fig. 2F and S3B**). Notably, LBs are also known to be marked by SQSTM-1 and autophagy markers including LC3 (19, 37, 38). Similar to canonical LBs, bigger perinuclear IBs of Syn-filaments partially co-localized with membranous organelles like mitochondria, (ER) and lysosome, and partly distorted their morphology and cellular distribution (**Fig. 2G**, yellow arrowheads **and S3C**).

The perinuclear non-filamentous SNCA(DM)-EGFP blobs, detected in paraformaldehyde fixed cells, were also stained by p129-Syn (**Fig. 2Ei and S3A**). SQSTM1 encircled these blobs irrespective of the occurrence of Syn-filaments (**Fig. 2H and S3D**, stick arrowheads). Perinuclear blobs stained by ubiquitin antibody were also prominent (**Fig. 2I and S3E**, arrowheads). When stained with γ-tubulin antibody, distinct perinuclear dots were noticed in uninduced cells (**Fig. 2Ji**, arrowhead). Perinuclear γ-tubulin was encircled by SQSTM1 upon induction of SNCA(DM)-EGFP and were observed even in presence of Syn-filaments (**Fig. 2Jii–iii and S3F**, stick arrowheads).

Thus, similar to the primary neurons, two distinct α-Synuclein IBs were also observed in mitotic cells. Switching misfolded proteins from aggresome to IPOD and *vice versa* is reported in literature but it always results in sequestration of the protein into one of the IBs (7). Deposition into co-existing IBs in the same cells was thus unique for α-Synuclein. Also, the second IB in this case, the LB-like perinuclear Syn-IBs, was externally seeded. Similar seeding-based propagation of α-Synuclein inclusions are reported in patient brains (39, 40). The question remained what could be the homeostatic benefits or drawbacks of having the seeded Syn-IBs while canonical aggresome-like perinuclear blobs were present in the same cells. Therefore, we next investigated the homeostatic contributions of the two independent IBs of α-Synuclein and compared them with the canonical QC-deposits.

### Syn-aggresomes and LB-like inclusions offer unique proteostatic opportunities

α-Synuclein is a known UPS-substrate. We found that p129-Syn degraded faster than the non-phosphorylated counterpart in cycloheximide (CHX) chase assay (**Fig. 3A**). Protein load of p129-Syn was increased after proteasome inhibitor MG132 treatment (**Fig. 3B**). Simultaneously, p129-Syn was dispersed throughout the cells rather than being restricted within the perinuclear blobs indicating that Ubiquitin-Proteasome System (UPS) works on substrate p129-Syn at this site (**Fig. 3B**, stick arrowhead). Further, encircling of the SNCA(DM)-EGFP blobs by SQSTM1 (**Fig. 2H**) suggested either degradation of p129-Syn by autophagy or degradation of the accumulated proteasomes at these aggresome-like IBs by STUB1 pathway (21). Microtubules contribute to the biogenesis of canonical aggresomes/JUNQs (12). SNCA(DM)-EGFP blobs disappeared within 30 min of Nocodazole treatment (**Fig. 3C** – above, stick arrowhead), a small molecule that disrupts microtubule assembly. Simultaneously, CHX chase assay revealed degradation of p129-Syn was delayed in nocodazole-treated cells (**Fig. 3C**, bottom). Thus, the perinuclear SNCA(DM)-EGFP blobs resembled canonical aggresomes serving a spontaneous cellular response to sequester degradable α-Synuclein. In presence of Syn-aggresomes, both α-Synuclein and p129-Syn never accumulated in excess in PFF-unseeded cells even after 6 days of continuous expression (**Fig. 3D**).

**Fig. 3:**
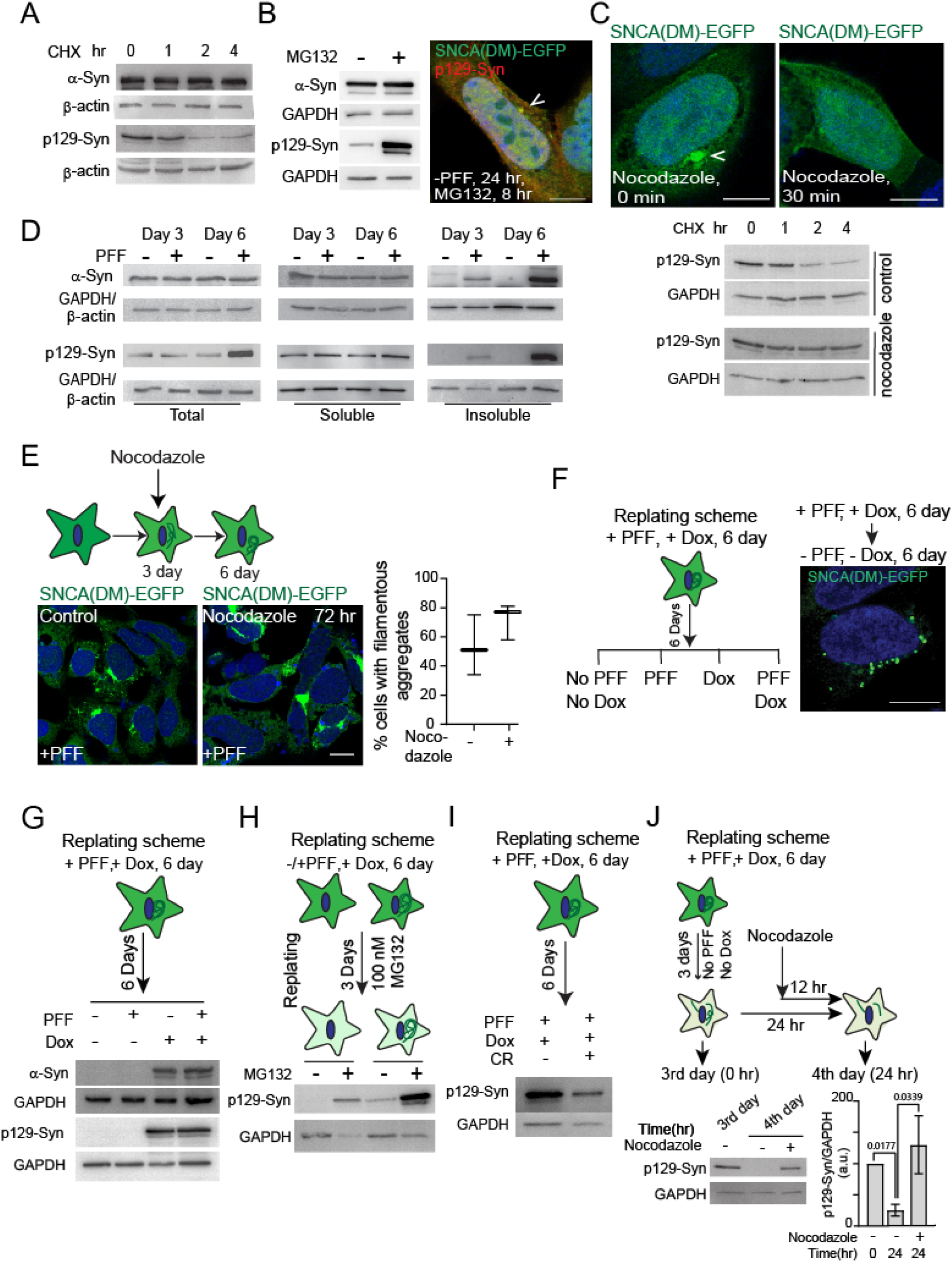
Syn-aggresomes and LB-like IBs offer unique homeostatic opportunities. **A**: 24 hr induced SNCA(DM)-EGFP cells treated with cycloheximide (CHX, 0.5 mM) for the indicated time points, total cell extract probed with anti α-Syn antibodies. α-Syn-detecting both phosphorylated and non-phosphorylated α-Synuclein and p129-Syn-detecting only phosphorylated α-Synuclein. β-actin as loading control. **B: Left**: 24 hr induced SNCA(DM)-EGFP cells treated with MG132 (5 µM) for 8 hr and total cell extract prepared from the cells immunoblotted for α-Syn and p129-Syn. GAPDH as loading control. **Right**: Same cells immunostained with p129-Syn antibody. Stick arrowheads indicate perinuclear blob. **C: Top**: 24 hr induced SNCA(DM)-EGFP cells, Nocodazole (0.1µg/ml, for 30 min) treated or untreated, were imaged. **Bottom**: Same cells were also treated with cycloheximide (CHX, 0.5 mM) together with nocodazole for indicated time points and immunoblotted for p129-Syn. GAPDH as loading control. **D:** SNCA(DM)-EGFP cells incubated with or without PFF for indicated days; total, soluble and insoluble extracts prepared as described in **Fig. S3G**, and probed for α-Syn and p129-Syn. GAPDH as loading control for total and soluble fractions, β-actin for insoluble fraction. **E: Top:** Experimental scheme of nocodazole (0.1μg/ml) treatments on 3^rd^ day for cytoskeleton destabilization and imaged on 6^th^ day. **Bottom** l**eft**: Microscopic image of control and nocodazole (0.1μg/ml) treated cells at 6^th^ day. **Bottom Right:** Number of SNCA(DM)-EGFP cells with filamentous IBs on 6^th^ day after 3 days of nocodazole treatment. Approx. 30 cells have been counted in each experiment. N = 3. **F: Left**: Re-plating experiment scheme. 6^th^ day filament containing cells were re-plated with or without doxycycline and PFF as indicated in the scheme. **Right**: Microscopic image of a re-plated cell. Re-plating conditions indicated. **G:** Experimental scheme - Extracts of cells after 6 day re-plating at indicated conditions, immunoblotted for α-Syn and p129-Syn. GAPDH as loading control. **H:** Experimental scheme - PFF untreated and treated 6^th^ day SNCA(DM)-EGFP cells were re-plated and 100 nM MG132 was added at every 24 hr for 3 days. 3^rd^ day cells collected; total extract probed for p129-Syn. GAPDH as loading control. **I:** Experimental scheme – Syn-filament containing cells were re-plated for 6 days with doxycycline and PFF ± Congo Red. Total cell extract probed for p129-Syn. GAPDH as loading control. **J: Top:** Experimental scheme: 3 days after re-plating, nocodazole (0.1μg/ml) was added 12 hr prior to sample collection at 4^th^ day. **Bottom Left:** Total cell extracts collected at indicated time points and probed for p129-Syn. **Bottom Right:** Graph represent the western blot quantification of 6 independent experiments. Students’s t-test. GAPDH as loading control. Scale bar – 10 µm. **Fig. 3B–C** – Paraformaldehyde fixation. **Fig. E-F** – Acetone:Methanol fixation.

In contrast to Syn-aggresomes, microtubule destabilization did not prevent perinuclear deposition of the Syn-filaments into LB-like IBs rather expedited its biogenesis (**Fig. 3E**) suggesting that the LBs observed in post-mortem PD brains are not the cytoskeleton-dependent canonical aggresomes. Interestingly, total α-Synuclein load also did not increase in PFF-treated 6^th^ day SNCA(DM)-EGFP cells although p129-Syn level was higher (**Fig. 3D**). Lysing the cells with 1% Triton-X 100 yielded soluble and insoluble fractions (**Fig. S3G**). α-Synuclein or p129-Syn load remained unchanged in the soluble fraction but both were increased in the insoluble fraction (**Fig. 3D**). The increase of α-Synuclein in the insoluble fraction blots was not justifiably proportionate to the large LB-like IBs observed in microscopy indicating ‘biochemical softness’ of these filamentous IBs. These blots further suggested that while a fraction of α-Synuclein was being degraded *via* Syn-aggresome, another fraction was increasingly phosphorylated, polymerized into Syn-filaments in presence of PFF, and turned sparingly insoluble. Remarkably, increase in p129-Syn level in total and insoluble fraction of 6^th^ day PFF-treated SNCA(DM)-EGFP cells was prevented in presence of Congo Red (**Fig. S3H**) that compromised polymerization efficiency of Syn-filaments (**Fig. S2G**).

Next, we designed a re-plating assay to investigate the stability of LB-like Syn-IBs (**Fig. 3F**). Sixth day PFF-treated cells with LB-like IBs were trypsinized and re-plated without PFF-seeds or doxycycline that induces synthesis of new SNCA(DM)-EGFP. The large LB-like IBs were first fragmented into multiple scattered dots (**Fig. 3F**). LB-like IB containing cells disappeared slowly while only few fragmented IB containing cells remain (**Fig. S3I**, left). On the other hand, cells with LB-like IBs remain in population if doxycycline and/or PFF-seeds were continued during re-plating (**Fig. S3I**, right). α-Synuclein and p129-Syn levels disappeared in western blots in absence of doxycycline and/or PFF-seeds during re-plating (**Fig. 3G**) while MG132 addition prevented the disappearance of p129-Syn band confirming UPS-mediated degradation (**Fig. 3H**). Intriguingly, treating the cells with Congo Red during re-plating resulted in reduction of p129-Syn even in presence of PFF and doxycycline (**Fig. 3I**) suggesting that impeding polymerization of fresh filaments was sufficient to disintegrate and degrade the LB-like IBs. In contrast, re-plating in presence of Nocodazole did not reduce the p129-Syn level (**Fig. 3J**) advocating a role of Syn-aggresomes in the degradation of p129-Syn accumulated into the LB-like IBs. Together, we conclude that perinuclear LB-like IBs facilitated aggresome-assisted opportunistic degradation of Syn-amyloids.

### Cells with perinuclear LB-like IBs are proteostasis-sensitive

Endogenous α-Synuclein accumulated in Syn-aggresomes without affecting primary neuron culture. PFF-untreated Syn-aggresome containing SNCA(DM)-EGFP cells also proliferated efficiently. As reported, biogenesis of LB-like IBs in primary neurons do not impact neuronal viability up to Day 14 in culture (19). Similarly, LB-containing primary neurons did not show significant LDH release up to 10 days of PFF incubation in our experiments (**Fig. 4A**). Also, cell viability remained unaffected despite the presence of LB-like IBs in 6^th^ day PFF-treated SNCA(DM)-EGFP cells (**Fig. 4B**). Moreover, Syn-filaments were asymmetrically partitioned into the daughter cells without compromising division (**Fig. 4Ci, S4A and Supplementary Video 2**) although infrequent nuclear entry of large LB-like IBs resulted in fragmentation of nuclei and cell detachment from the plate surface in some cases (**Fig. 4D, S4B and supplementary video 3**).

**Fig. 4:**
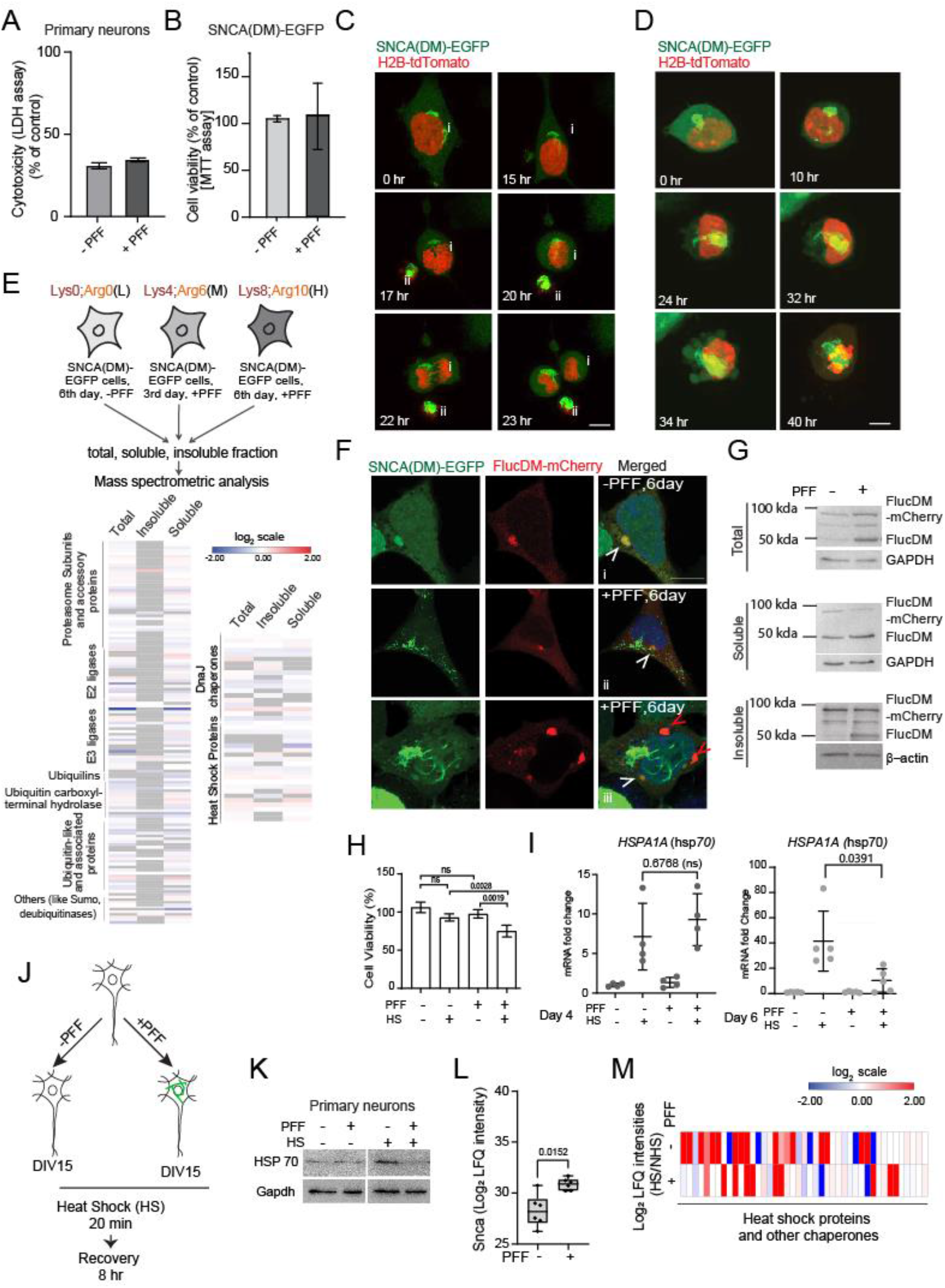
Cellular homeostasis is near saturation in LB-like IB containing cells. **A**: Cytotoxicity estimated by LDH release assay at DIV15 after 10 days without and with 10 days PFF incubation in primary neurons. LDH release in a fresh cell lysate taken as positive control (100%). The percentage of LDH release relative to positive control plotted. Error bars indicate SD. N=4. **B:** Cell viability of 6^th^ day PFF treated SNCA(DM)-EGFP cells as estimated by MTT assay. Error bars indicate SD. N=3. **C:** Different stages of mitotic division of LB-like IBs containing cell as captured by live cell microscopy. The cells were transfected with H2B-tdTomato as nuclear marker (by virtue of its affinity to chromatin (41) at 3^rd^ day of PFF incubation. Live imaging started on 4^th^ day of PFF incubation for 72 hrs. (**i**). A detached floating cell shown in (**ii**). **D:** Detachment of LB-like IB containing cell from plate surface as captured by live cell microscopy. **E: Top:** Experimental scheme. **Bottom:** Heatmap showing fold-change (SNCA(DM)-EGFP, 6^th^ day; Ratio: +PFF/-PFF) distribution of PQC-proteins in total, soluble and insoluble fractions. **F:** Localization of FlucDM-mCherry in 6^th^ day PFF treated and untreated SNCA(DM)-EGFP cells. Cells were transfected with FlucDM-mCherry on 5^th^ day of incubation and fixed at day 6 with paraformaldehyde. **G:** Total, soluble and insoluble extracts were prepared from cells shown in **4F** and probed using Luciferase antibody. ∼100 kDa band – full length FlucDM-mCherry; 50 kDa band – cleaved FlucDM. GAPDH/ β-actin as loading control. **H:** MTT assay - 6^th^ day SNCA(DM)-EGFP cells (-/+) PFF incubation after 2 hr heat shock at 42°C. N=5, Student’s *t*-test. **I:** 4^th^ day and 6^th^ day (-/+) PFF SNCA(DM)-EGFP cells treated with HS – heat stress at 42°C for 2 hr. RT-PCR was performed for *HSPA1A*. Fold change compared to GAPDH control plotted. Welch *t*-test. **J:** Experimental Scheme - Untreated and PFF treated primary neurons were subjected to heat shock (HS) at 42 °C for 20 minutes and followed by recovery for 8 hr. Cells were collected for western blot and mass spectrometry analysis. **K:** Total extract prepared from primary neurons with heat shock (HS) and non-heat shock (NHS) as shown in **Fig. 4J** were probed with HSP70 antibody. Gapdh as loading control. **L:** Log2 LFQ intensities for α-Synuclein from unseeded and PFF treated primary neurons as quantified by mass spectrometry. The experimental scheme shown in **Fig. 4J**. Student’s t-test. **M:** Heat map showing (HS/NHS) ratio of LFQ intensities from untreated and PFF treated primary neurons as quantified by mass spectrometry. Experimental scheme shown in **Fig. 4J**. N=3. Scale bar – 10 µm.

SILAC-based quantitative mass spectrometry of the total, soluble and insoluble fractions of SNCA(DM)-EGFP cells revealed almost unperturbed proteome in presence of both Syn-IBs (**Fig. S4C–D, Table S1–S3**). Despite the recruitment of multiple PQC factors into Syn-aggresomes and large perinuclear LB-like IBs as observed in microscopy (**Fig. 2F, H–I**), chaperones and degradation machinery components were not increased in 6^th^ day PFF treated SNCA(DM)-EGFP cells (**Fig. 4E, Table S2**). This suggested that the recruitment of PQC-factors onto the LB-like IBs was transient, not stable enough to be precipitated into the insoluble fraction. Mitochondrial proteins were non-significantly increased in all the proteome fractions suggesting a possible adjustment in mitochondrial biology (**Fig. S4C, Table S1**). Nuclear proteins were slightly elevated in soluble fraction while decreased in insoluble fraction (**Fig. S4C**).

To test whether transient repurposing of PQC-factors to the Syn-IBs could trigger proteostasis-insufficiency for other metastable proteins, we overexpressed misfolding-prone FlucDM-mCherry (42) in 6^th^ day SNCA(DM)-EGFP cells. FlucDM-mCherry was deposited into the Syn-aggresomes in absence of PFF suggesting that the spontaneous cellular response of directing metastable proteins into the perinuclear QC-sites remained active in SNCA(DM)-EGFP cells (**Fig. 4Fi**, arrowhead). Excess FlucDM-mCherry did not accumulate in these cells confirming that co-presence of SNCA(DM)-EGFP in the aggresomes did not compromise its turnover (**Fig. 4G**). Remarkably, in addition to partial co-localization with Syn-aggresomes and LB-like IBs (**Fig. 4Fii**, white arrowheads), FlucDM-mCherry appeared in additional cytoplasmic IBs in 6^th^ day PFF-treated SNCA(DM)-EGFP cells (**Fig. 4Fiii**, red arrowheads). Protein-load of FlucDM-mCherry was also increased in total and insoluble fractions (**Fig. 4G**). These results suggested that the mechanism of depositing misfolding-prone proteins into canonical QC-deposits was saturated in presence of LB-like IBs. Intriguingly, separate IBs of FlucDM-mCherry were absent in cells with small Syn-filaments (**Fig. 4Fii**) indicating that over-saturation of cellular-PQC was manifested only after maturation of large perinuclear LB-like IBs.

To further investigate the proteostatic-response of LB-like IB containing cells against widespread proteome misfolding stress, we incubated the 4^th^ and 6^th^ day SNCA(DM)-EGFP cells at 42°C for 2 hr. Viability of the PFF-treated cells was significantly reduced after thermal stress on 6^th^ day (**Fig. 4H**). Transcription upregulation of stress chaperone Hsp70 was also deregulated in heat stressed 6^th^ PFF-treated cells substantiating their inability to trigger the synthesis of fresh PQC-factors (**Fig. 4I**). In contrast, Hsp70-transcription remained normal in 4^th^ day cells when only small peripheral Syn-filaments were observed suggesting that transcription deregulation of Hsp70 was correlated with the appearance of LB-like Syn-IBs (**Fig. 4I**). To note, heat stress did not perturb the perinuclear location of any of the Syn-IBs (**Fig. S4E**).

Likewise the 6^th^ day SNCA(DM)-EGFP cells, Hsp70 protein level was not triggered by heat stress in PFF-treated primary neurons (**Fig. 4J–K**). Quantitative mass spectrometry indicated increased accumulation of α-Synuclein after PFF-treatment (**Fig. 4L**). Chaperone levels were alternatively upregulated after heat stress in the same PFF-treated neurons suggesting gross defects in the stress response pathways (**Fig. 4M and Table S4**).

### Perinuclear LB-like IBs trigger nuclear lamina deformities

Syn-filaments accumulate into LB-like IBs at the perinucleus (**Fig. 1Bii and 2C**). Occasional intra-nuclear localization of Syn-filaments and nuclear fragmentation was also observed in live cell microscopy (**Fig. 4D**). Transcription is a nucleus-mediated cellular function and was deregulated in cells with perinuclear LB-like IBs (**Fig. 4I**). These observations prompted us to investigate the nuclear shape, size and integrity in detail.

Staining with nuclear envelope (NE) protein Lamin B1 (LmnB1) revealed increased nuclear lamina discontinuity in cells with LB-like IBs (**Fig. 5A–B**, arrows). Often, LB-like IBs overlapped LmnB1 staining and lamina disintegrities were observed both at the and far to the overlap sites (**Fig. 5Bii–iii**). Co-staining with LmnB1 was observed at the core (**Fig. 5Bi–ii**) and also at the outer periphery of the LB-like IBs (**Fig. 5Biv**). Symmetric distribution of LmnB1 across the NE was lost with accumulation of LmnB1 fluorescence at the overlap sites with LB-like IBs (**Fig. 5C**) suggesting mechanical squeezing the nuclear lamina. Smaller Syn-filaments, both perinuclear and non-perinuclear, remained unstained by LmnB1 (**Fig. S5A**). Rarely, Syn-filaments co-stained with LmnB1 were found outside the intact NE (**Fig. 5Bi**) suggesting re-establishment of nuclear lamina by disposing of the injured part. Intriguingly, lamina injuries in presence of LB-like IBs did not alter LmnB1 protein level (**Fig. S5B**) although resulted in slight but significant drop in nuclear volume (**Fig. S5C**).

**Fig. 5:**
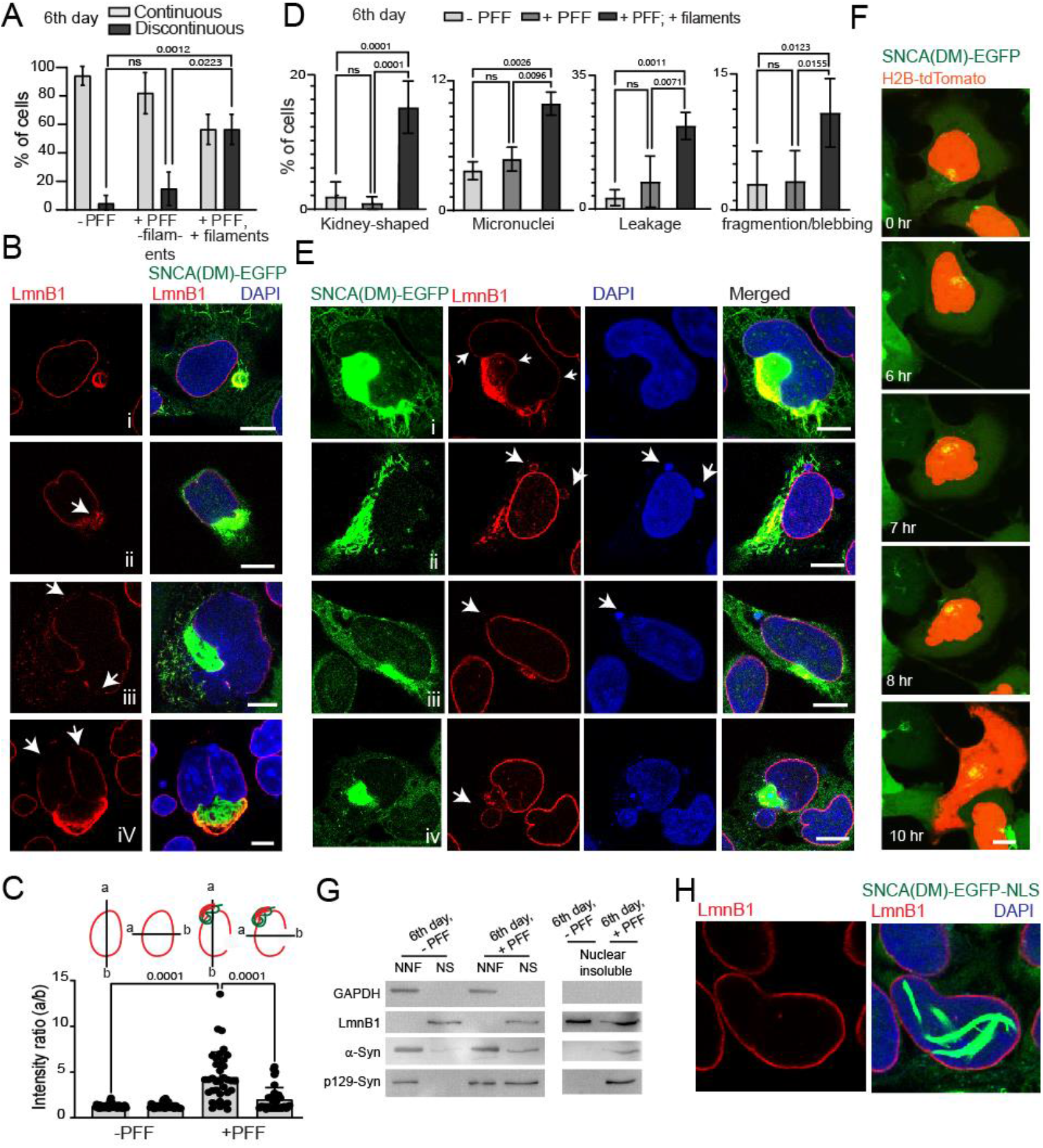
Perinuclear LB-like IBs trigger nuclear lamina deformities. **A-B**: 6^th^ day PFF treated SNCA(DM)-EGFP cells immunostained for LmnB1. Percentage of cells with continuous or discontinuous lamina as per LmnB1 staining plotted (**A**). Error bars – SDs. N=5. Approx. 100 cells counted in each biological repeat. Student’s t-test. Representative images shown (**B**). Arrows indicate irregularities in LmnB1 staining. **C:** Ratio of LmnB1 intensity distribution on NE calculated as indicated in 6^th^ day PFF treated and untreated SNCA(DM)-EGFP cells. Error bars – SDs. N=3. Student’s t-test. **D-E:** 6^th^ day PFF treated SNCA(DM)-EGFP cells immunostained for LmnB1. Deformities of nuclear morphology and appendages as per LmnB1 and DAPI staining plotted (**D**). Error bars – SDs. N=5. Approx. 100 cells were counted in each biological repeat. Student’s t-test. Representative images shown (**E**). Arrows indicate nuclear deformity sites. **F:** Live cell microscopy images showing cellular localization of SNCA(DM)-EGFP and H2B-tdTomato in Syn-filament containing cells. Cells transfected with H2B-tdTomato on 3rd day of PFF incubation. Live imaging started on 4^th^ day of PFF incubation for 72 hrs. **G:** Non-Nuclear (NNF), Nuclear soluble (NS) and Nuclear insoluble fractions prepared from 6^th^ day SNCA(DM)-EGFP (+/-PFF) cells and probed for LmnB1, GAPDH, α-Syn and p129-Syn. Fractionation procedure shown in **Fig-S5E**. **H:** 6^th^ day PFF treated SNCA(DM)-EGFP-NLS cells were fixed and immunostained with LmnB1 antibody **Fig. 5 B, E, H** -Acetone:Methanol fixation. **Fig. 5F** – Live cell imaging.

We categorized the nuclear deformities in these cells according to DAPI and LmnB1 staining (**Fig. 5D–E**, arrowheads). Kidney-shaped nuclei were observed in ∼ 15-20 % of the cells with LB-like IBs confined within the NE-invaginations (**Fig. 5Ei**). LmnB1 positivity of these perinuclear IBs indicated local lamina injury at the overlap sites. As seen in laminopathies (43–45), nuclear hernia-like blobs were captured in live cell microscopy (**Fig. S5D and Supplementary video 4**). Extranuclear DAPI blobs confined within LmnB1 boundary were termed as micronuclei (**Fig. 5Eii**) while LmnB1-negative DAPI blobs close to the lamina discontinuity sites were considered as leakages (**Fig. 5Eiii**). Large spherical LmnB1-positive DAPI-lobules were found in more than 10% of the cells with LB-like IBs and were classified as fragmented or blebbed nuclei (**Fig. 5D and 5Eiv**). Similar nuclear blobs and lobules were observed in the detaching cells in our live cell microscopy experiments suggesting necrosis-like cell death (**Fig. 4D and Supplementary video 3**). Nuclear rupture in few cells with LB-like IBs resulted in leakage of H2B-tdTomato into the cytoplasm which was not observed in absence of the perinuclear filamentous IBs (**Fig. 5F and Supplementary video 5**).

Next, we prepared nuclear and non-nuclear fractions as depicted in **Fig. S5E**. Both phosphorylated and non-phosphorylated forms of α-Synuclein increased in the soluble and insoluble nuclear fractions of 6^th^ day PFF treated SNCA(DM)-EGFP cells (**Fig. 5G**). LmnB1 was mostly present in the insoluble nuclear fraction. Appearance of α-Synuclein bands in the same insoluble fraction of PFF-treated cells indicated association between insoluble LmnB1 and LB-like IBs (**Fig. 5G**). Mitochondria and ER were found close to the nucleus and partially co-localized with LB-like IBs (**Fig. 1D and 2H**). Non-significant but consistent increase of mitochondrial and ER proteins in the nuclear fractions of the PFF treated cells indicated entry of a small fraction of organelle proteins through the leaky NE (**Fig. S5F–G, Table S5-S6**).

Notably, lamina injuries were repaired with the disappearance of LB-like IBs upon withdrawal of PFF or doxycycline during re-plating (**Fig. S5H**). We also tagged nuclear localization signal at the C-terminal of SNCA(DM)-EGFP. SNCA(DM)-EGFP-NLS deposited into intranuclear Syn-filaments instead of perinuclear LB-like IBs (**Fig. 5H**). Remarkably, lamina remained intact despite having large intranuclear Syn-filaments suggesting that the lamina damaging interactions of LB-like Syn-IBs were confined at the perinucleus.

Co-staining of perinuclear LB-like IBs with LmnB1, lamina discontinuity, kidney-shaped nucleus etc. were consistent after 10 days of PFF-incubation in DIV15 primary neurons (**Fig. 6Aii–iii and S5I**). In contrast, nuclear lamina remained unperturbed in neurons with smaller peripheral Syn-filaments and in absence of PFF treatment (**Fig. 6Ai and S5I**). Disruption of lamina boundary and mixing of nucleocytoplasmic components often trigger DNA-damage (46). Multiple nuclear γ-H2AX foci confirmed DNA-damage in neurons perinuclear LB-like IBs (**Fig. 6Bii and S5J**) while neighbouring neurons with small peripheral filaments displayed 1-2 faintly stained γ-H2AX foci (**Fig. 6Bi and S5J**). Next, we verified if the lamina injuries were consistent in a more contextual pathophysiological system. We stained post-mortem thalamus sections of a Parkinson’s Disease (PD) patient (BioChain Institute Inc., USA Cat# T2236079Par) and a control (BioChain Institute Inc., USA Cat#T2234079) with p129-Syn and LmnB1 antibodies. We observed extensive filamentous staining by p129-Syn antibodies in the patient brain suggesting Lewy neurites and Lewy Bodies (47) while the control was relatively unstained (**Fig. 6C**). Extensive staining of p129-Syn at the perinucleus was co-related with mechanically deformed nucleus (**inset, Fig. 6Ci**) while small filaments at the periphery was associated with normal nuclear morphology (**inset, Fig. 6Cii**). The mechanical deformities of the nuclei in the patient brain section were confirmed by LmnB1 staining which indicated widespread lamina injuries compared to the control brain (**Fig. 6D–E**).

**Fig. 6:**
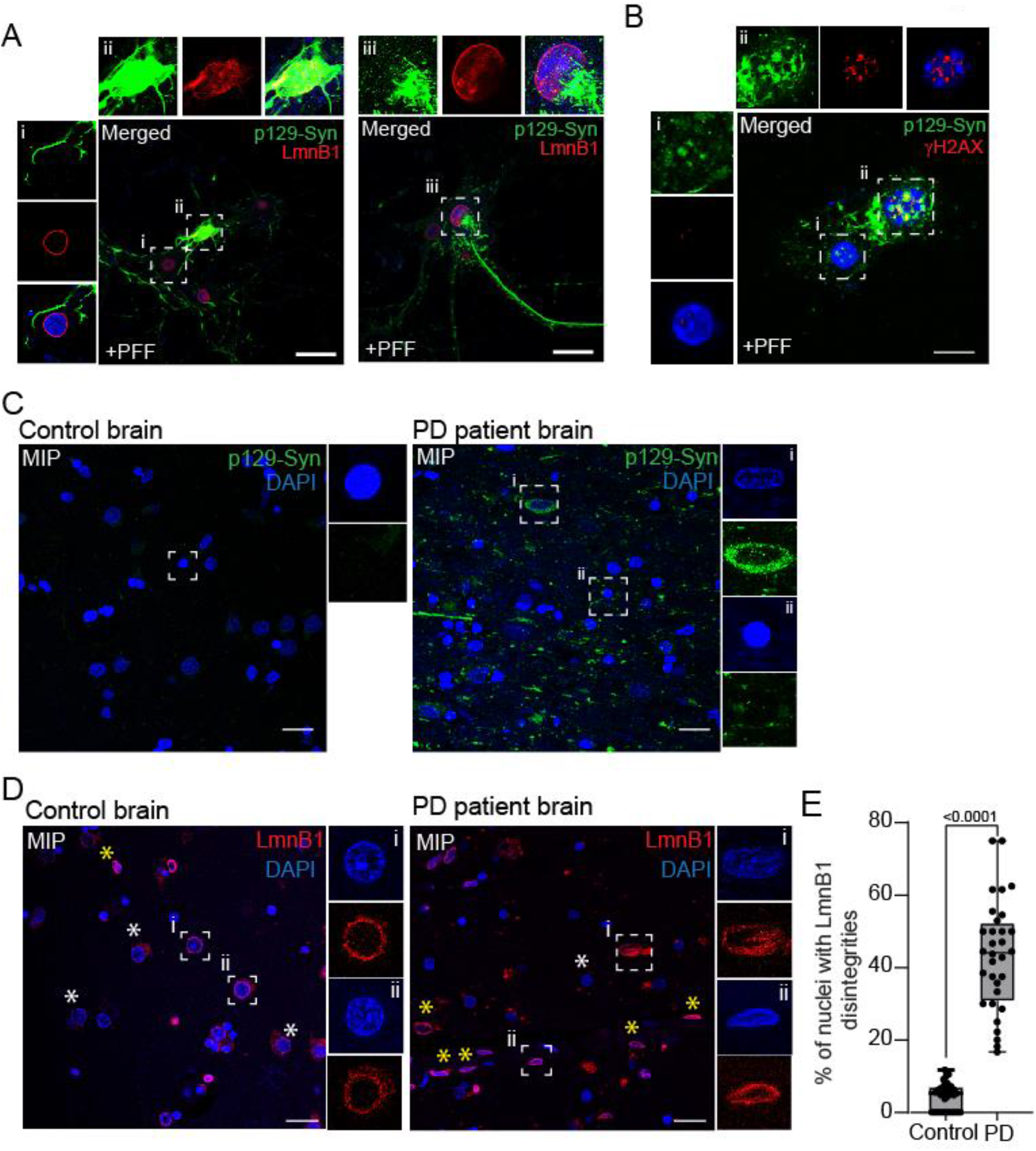
LBs trigger nuclear lamina deformities. **A**: Primary neurons at 15 DIV, 10 days post PFF treatment fixed and immunostained with p129-Syn (P-syn/81A) and LmnB1. Insets: zoomed images of the indicated locations. Scale bar-10 µm. **B:** Primary neurons at 15 DIV, 10 days post PFF treatment fixed and immunostained with p129-Syn (MJF-R13 (8-8)) and γH2AX antibody. Insets: zoomed images of the indicated locations. Scale bar-10 µm. **C:** MIP of a non-neurological disease (control) and Parkinson’s disease patients’ brain sections were stained with p129-Syn (MJF-R13 (8-8)) antibody. Scale bar-20 µm. insets: zoomed images of box-indicated locations. **D:** MIP of a non-neurological disease (control) and Parkinson’s disease (PD) patients’ brain sections were stained with LmnB1 antibody. insets: zoomed images of box-indicated locations. Scale bar-20 µm. White asterisk – representative nuclei with unperturbed lamina; Yellow asterisk – nuclei with lamina disintegrities. More representative microscopy fields of brain sections are shown in **Fig. S6**. **E:** The percentage of nuclei with lamina disintegrities among total LmnB1 positive nuclei per field of control brain and Parkinson’s disease (PD) patients’ brain sections were plotted. N = 1, Total 49 field of control and 44 field of PD brain sections of dimensions 185×185 µm were counted. Average 12-15 LmnB1 positive nuclei per field. **Fig. 6A–B** – Acetone:Methanol fixation. **Fig. 6C–D** – formalin fixed/paraffin embedded brain sections.

### Lamina-injuries by misbalanced cytoskeleton-forces responsible for proteostasis sensitivity in presence of LB-like IBs

Nuclear lamina represents the interior meshwork of the nuclear envelope (NE) below the lipid bilayers. Given the size of the perinuclear LB-like IBs was far beyond the channel capacity of nuclear pore complexes (48), we asked how these large IBs crossed the lipid bilayer, overlapped with and injured the lamina. One possibility was aberrant association of NE-lipids with LB-like IBs due to intrinsic lipid-affinity of the latter (39, 49, 50). Indeed, LB-like IBs colocalized with membranous organelles at the perinuclear space (**Fig. S1D–E and 2G**). Also, LB-like IBs were positive for lipid-staining dye DiIC_16_ (1,1’-Dihexadecyl-3,3,3’,3’-Tetramethylindocarbocyanine Perchlorate) at the perinucleus (**Fig. S7A**, yellow arrow). These experiments indicated that the lipid affinity of the LB-like IBs could facilitate its association with NE but could not explain the underlying cause of lamina disintegrities.

Perinuclear cytoskeleton connects to nucleoskeleton via LINC (Linker of Nucleoskeleton and Cytoskeleton) complex proteins and determine the intracellular positioning and integrity of nucleus (51, 52). Aberrant cytoskeleton forces at the perinucleus can trigger lamina-injuries. In steady-state cells, cytosolic motor proteins transport the cargos towards and away from the nucleus. For example, dynein carries peripheral misfolded proteins along the microtubules for deposition into aggresomes at the perinucleus (53).

On the other hand, dynein-force also pulls the NE towards the MTOC during mitosis. This tears the lamina elsewhere (54). Dynein pulling forces aggravate NE-injuries even in interphase cells if the lamina is weak (54, 55). Actin-cytoskeleton also confines and mechanically squeezes the nucleus at interphase to rupture lamina (45). Intriguingly, LB-like IBs and MTOCs were often present together within the NE-invaginations of the kidney-shaped SNCA(DM)-EGFP cells (**Fig. 7A**). This is very similar to MTOC organization in prometaphase cells (54) and suggested that altered mechanical forces contributed by perinuclear cytoskeleton could be responsible for the NE-deformities observed in presence of LB-like IBs. To verify, we imaged neurons with LB-like IBs for cytoskeleton proteins.

**Fig. 7:**
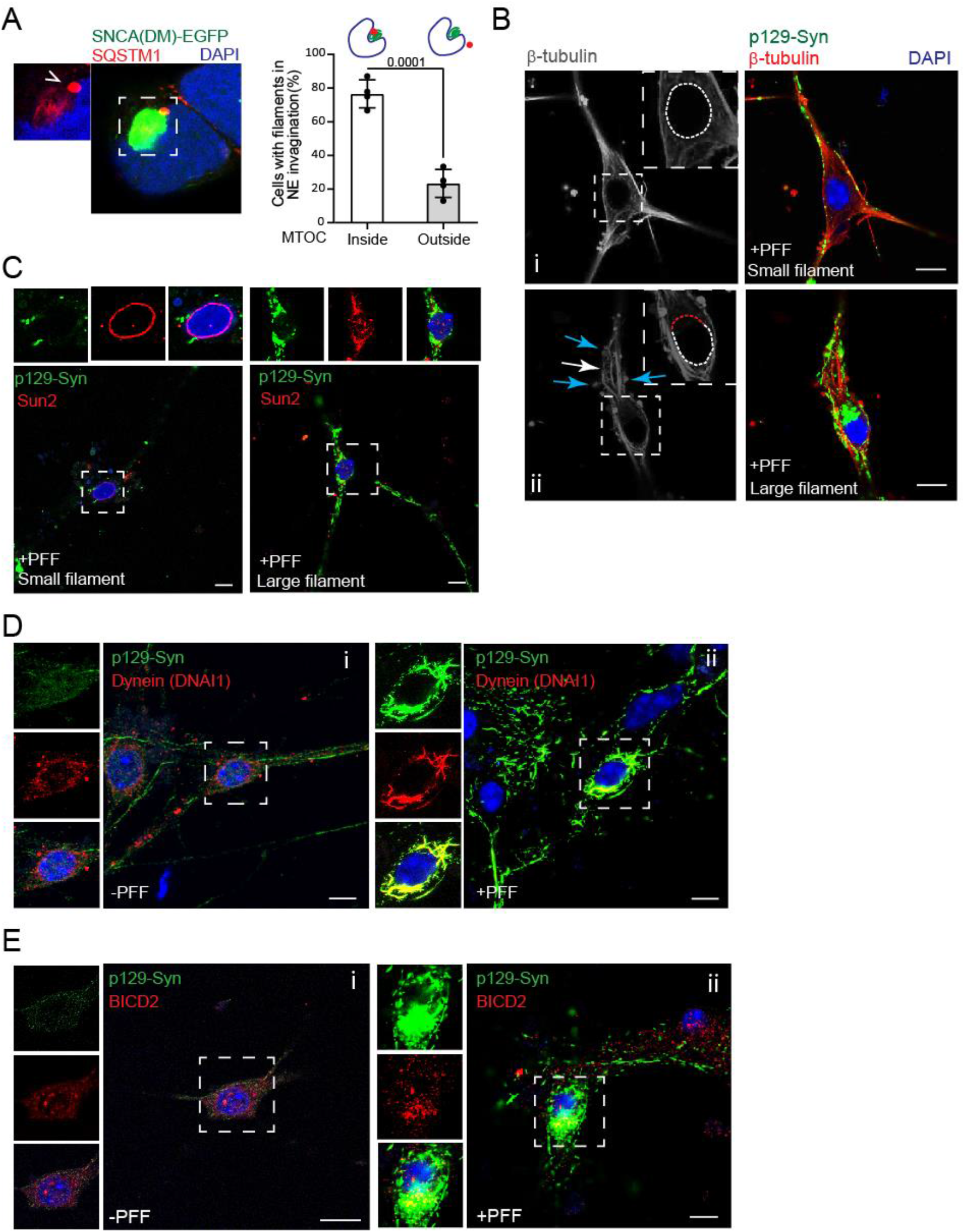
Perinuclear cytoskeleton in presence of LB-like IBs. **A**: Left: 6^th^ day PFF treated SNCA (DM)-EGFP cells stained with SQSTM1 antibody indicating aggresome location (Stick arrowhead). **Right:** No. of cells with aggresomes inside kidney shaped grooves counted and plotted. Error bars – SDs. N=3. Students t-test. **B:** Primary neurons at 15 DIV, 10 days post PFF treatment, fixed and immunostained with p129-Syn (P-syn/81A) and β-tubulin antibody. Red and white dotted circle indicate tubulin meshwork at nucleocytoplasmic boundary. White and blue arrows indicate bundles and tubulin blobs respectively. Insets – zoomed images of the indicated locations. **C:** Primary neurons at 15 DIV, 10 days post PFF treatment, fixed and immunostained with p129-Syn (P-syn/81A) and Sun2. Insets – zoomed images of the indicated locations. **D:** Primary neurons at 15 DIV, 10 days post (-/+) PFF treatment, fixed and immunostained with p129-Syn (P-syn/81A) and Dynein (DNAI1). Insets – zoomed images of the indicated locations. **E:** Primary neurons at 15 DIV, 10 days post (-/+) PFF treatment, fixed and immunostained with p129-Syn (P-syn/81A) and Anti-BICD2. Insets – zoomed images of the indicated locations. Scale bar-10 µm. All the cells shown in images were fixed with Acetone:Methanol.

The smooth microtubule meshwork remained unperturbed in neurons with peripheral Syn-filaments (**Fig. 7Bi and S7B**) but turned into bundles and blobs in presence of the perinuclear LB-like IBs (**Fig. 7Bii and S7B**, white and blue arrows). Further, the uniformity of the nuclear boundary disappeared close to the LB-like IBs (**Fig. 7Bii and S7Bii**, insets, red dotted nuclear boundary). Distribution of LINC complex protein SUN2 across the NE was also distorted as LB-like IBs encaged the nucleus suggesting perturbation of cytoskeleton-nucleoskeleton connections (**Fig. 7C and S7C**). Microtubule imparts pulling forces on the NE through dynein motors through the uniformly distributed LINC complex proteins on the NE (55). Cytosolic dynein was observed throughout the cell body and in the projections of DIV14 primary neurons in absence of PFF (**Fig. 7Di and S7D**). Uniform distribution of this motor protein was also noticed surrounding the nuclear boundary. This distribution symmetry was remarkably distorted in presence of PFF where cytosolic dynein motors were extensively positioned on the large filamentous Syn-IBs at the perinucleus (**Fig. 7Dii and S7D**). This suggested a deregulated dynein motor force could be working on the nucleoskeleton. Cytosolic cargo adapter protein BICD2 (Bicaudal D Homolog 2) is implicated in dynein-mediated motility of cargo along the microtubules. Lipid–rich membranous vesicles are common cargos for BICD2 (56, 57). Further, LINC-complex protein Nesprin-2 recruits dynein on NE via BICD2 (58, 59). Remarkably, while BICD2 was distributed throughout the cell body including the nuclear surrounding of control primary neurons (**Fig. 7Ei and S7E**), it co-localized massively with the perinuclear LB-like IBs in the 10^th^ day PFF-treated neurons (**Fig. 7Eii and S7E**). This suggested that BICD2-mediated dynein recruitment on the perinuclear LB-like IBs could be due to the misidentification of the IBs as lipid-rich membrane-wrapped cargos by BICD2.

Similar to primary neurons, microtubule organization was also destabilized at the periphery of LB-like IBs in Hek293T cells (**Fig. 8A**) while actin cytoskeleton remained comparatively unperturbed (**Fig. S8A**). The smooth β-Tubulin staining marking the nucleocytoplasmic boundary in control cells disappeared near the large filamentous IBs (**Fig. 8A**, red dotted nuclear boundary). Fragmented microtubule meshwork was also noticed at the cell-peripheral (**Fig. 8A**, blue arrow). LINC complex proteins Sun2 and Nesprin-2 were partially co-localized with perinuclear LB-like IBs and distributed as asymmetric clusters on the NE (**Fig. 8B and S8B**). Dynein was majorly deposited into a perinuclear blob in control cells, likely to be the aggresomes on MTOC (54), with faint staining surrounding the nucleus (**Fig. 8C**). BICD2 was also marking the nuclear boundary (**Fig. 8D**). Like primary neurons, both dynein and BICD2 re-localized onto the perinuclear LB-like IBs after PFF-treatment (**Fig. 8C–D**).

**Fig. 8:**
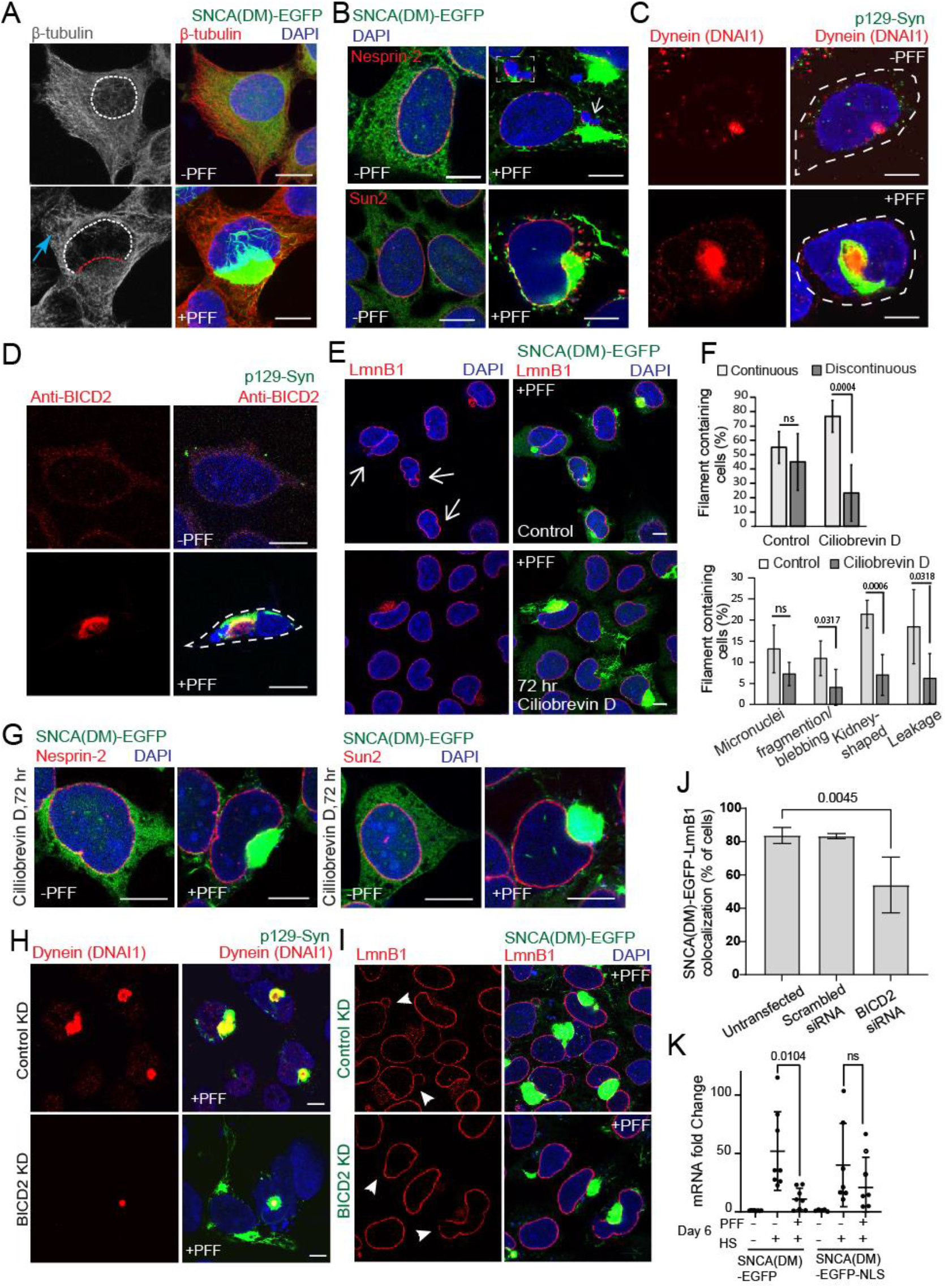
Altered cytoskeleton-forces in LB-like IB containing cells trigger NE-injuries. **A**: Maximum Intensity projection (MIP) of microscopic images of 6^th^ day (-/+) PFF treated SNCA(DM)-EGFP cells stained with β-tubulin antibody. Red and white dotted circle indicate tubulin meshwor at nucleocytoplasmic boundary. Blue arow indicate tubulin meshwork in peripheral region. **B:** Microscopic images of 6^th^ day (-/+) PFF treated SNCA(DM)-EGFP cells immunostained with Nesprin-2 and Sun2 antibodies. Insets – zoomed images of arrow indicated sites. **C:** Microscopic images of 6^th^ day (-/+) PFF treated SNCA(DM) cells immunostained with p129-Syn (P-syn/81A) and Dynein (DNAI1). Dotted line indicates cell boundary. **D:** Microscopic images of 6^th^ day (-/+) PFF treated SNCA(DM) cells immunostained with p129-Syn (P-syn/81A) and anti-BICD2. Dotted line indicates cell boundary. **E:** 6th day PFF treated SNCA (DM)-EGFP cells immunostained for LmnB1 after 72 hr Ciliobrevin D treatment (5 µM). **F:** Quantification of deformities in nuclear morphology as per LmnB1 staining on 6^th^ day after 72 hr Ciliobrevin D treatment (5 µM). Control – DMSO-treated cells. Error bars – SDs. N=5. Approx. 60 cells were counted in each experiment. Student’s t-test. **G:** Microscopic images of 6^th^ day (-/+) PFF treated SNCA(DM)-EGFP cells after 72hr of Ciliobrevin D treatment, immunostained with Nesprin-2 and Sun2 antibodies. **H:** 6th day PFF treated SNCA (DM) cells immunostained with p129-Syn (P-syn/81A) and Dynein (DNAI1) after 72 hr BICD2 knockdown (KD). **I:** 6th day, with and without PFF treatment, SNCA (DM)-EGFP cells immunostained with LmnB1 after 72 hr BICD2 knockdown (KD). White arrows indicate lamina gaps. **J:** Quantification of LmnB1 association with Syn-filaments on 6^th^ day as per LmnB1 staining after 72 hr BICD2 knockdown. Error bars – SDs. N=4. Approx. 90-100 cells were counted in each experiment. Student’s t-test. **K:** 6^th^ day (-/+) PFF treated SNCA(DM)-EGFP and SNCA(DM)-EGFP-NLS cells subjected to heat stress at 42°C for 2 hr. RT-PCR was performed for *HSPA1A*. N=8 SNCA(DM)-EGFP, N=7 SNCA(DM)-EGFP-NLS. Welch’s t-test. **Fig. 7 D, I** - Paraformaldehyde fixation. **Fig. 7 A-C, E, G, J** - Acetone:Methanol fixation. Scale bar – 10 µm

Functional dynein motors are important for aggresome biogenesis while non-functional dynein trigger a BAG1-mediated compensatory mechanism to target the misfolded proteins to proteasome (53), (60). Impairing of dynein motor by the ATPase domain inhibitor Ciliobrevin D (61) was sufficient to destabilize the Syn-aggresomes (**Fig. S8C**) but was unable to prevent biogenesis of the LB-like IBs (**Fig. 8E**). Remarkably, lamina discontinuity, NE-invaginations, and leakages were significantly reduced in the same Ciliobrevin D treated cells (**Fig. 8F**). Ciliobrevin D also partially restored the symmetric distribution of LINC complex proteins (**Fig. 8G and S8D**) indicating that the destabilized mechanical force on the nucleoskeleton was relatively neutralized after stopping the dynein motor. BICD2 knockouts trigger lamina disintegrities (58, 62). Similarly, siRNA mediated BICD2 knockdown in our cells also resulted in small discontinuities in lamina (**Fig. S8E**, arrows). At the same time, knockdown of BICD2 resulted in prominent reduction in the recruitment of dynein to the perinuclear LB-like IBs (**Fig. 8H**). Colocalization between large perinuclear LB-like IBs and LmnB1 was also significantly reduced (**Fig. 8I–J**). Intriguingly, delocalization of LB-like IBs from perinucleus by tagging SNCA(DM)-EGFP with C-terminal NLS resulted in intranuclear Syn-filaments. Despite being large in size and encompassing the whole nucleus, these intranuclear filaments did not co-localize with DAPI staining (**Fig. 5H**). Upregulation of stress chaperones was partially rescued in these cells after heat stress (**Fig. 8K**) suggesting that the transcriptional deregulations was associated with the lamina injuries caused by the perinuclear positioning of the LB-like IBs.

Lamins can counteract dynein pulling forces on the NE (51). Overexpression of LmnB1 can also increase lamina stiffness (63). LmnB1-mCherry overexpression in 6^th^ day PFF treated SNCA(DM)-EGFP cells resulted in multi-compartmentalized nucleus irrespective of the occurrence of Syn-filaments (**Fig. S8E**). Strikingly, lamina discontinuity, micronuclei, and nuclear leakages disappeared after LmnB1-mCherry overexpression despite partial co-localization of the large Syn-IBs with LmnB1 in the deformed nuclei (**Fig. S8F–H**).

Taken together, we propose a model explaining the aberrant cytoskeleton force disintegrating the nucleoskeleton (**Fig. 9A**). BICD2-mediated dynein recruitment on large perinuclear LB-like IBs destabilizes the local microtubule tension on the nucleoskeleton. The misbalanced dynein pulling forces squeeze and disintegrate the nucleoskeleton along with the lamina. Thereby, a part of lamina along with LINC complex proteins aberrantly associate with large perinuclear LB-like Ibs. Pharmacological inhibition of dynein motor or preventing dynein misrecruitment on LBs by knocking down BICD2 relaxes the deregulated tension on NE and re-establishes nucleoskeleton symmetry.

**Fig. 9A.**
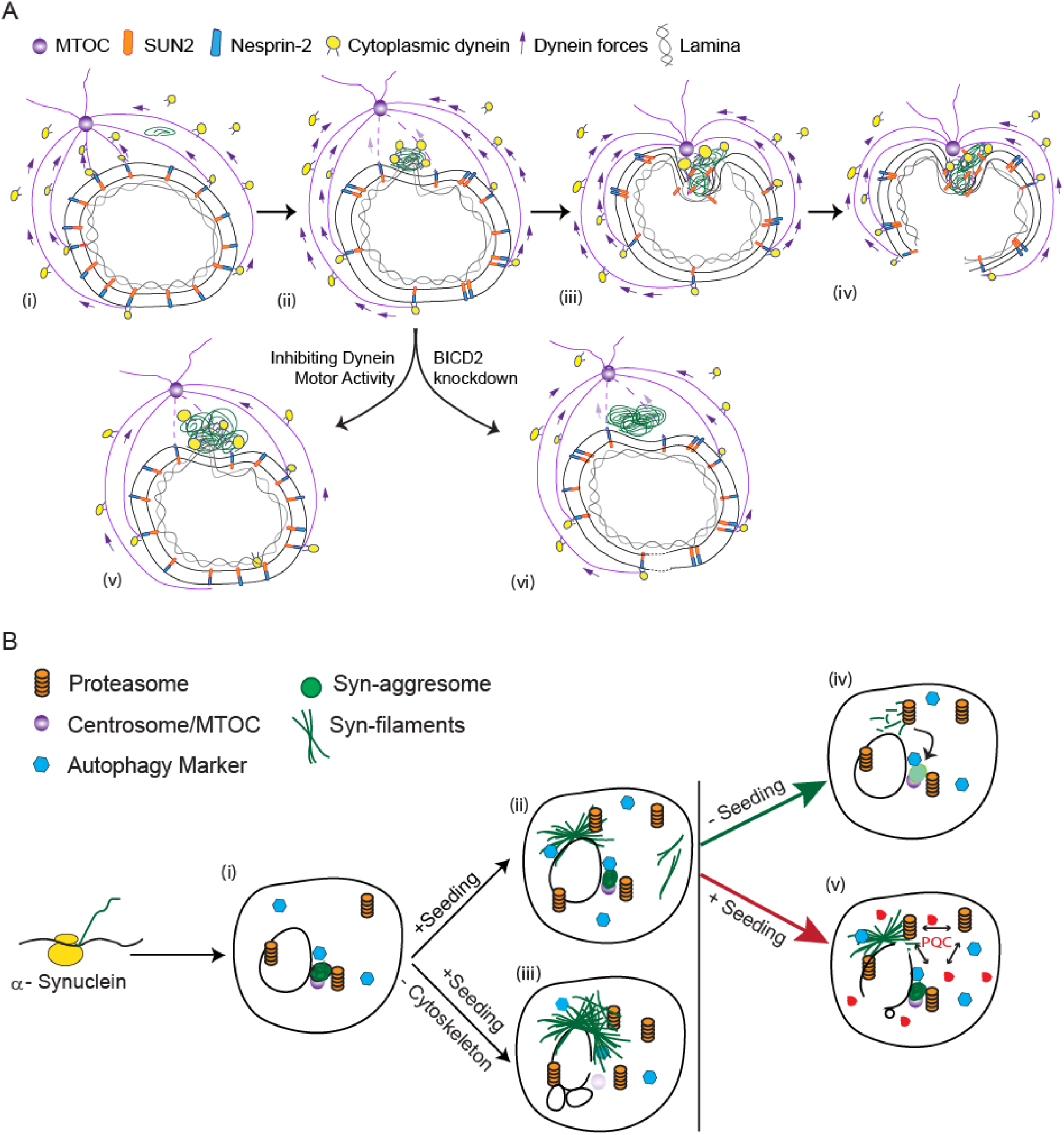
Mechanistic model for LB-induced nuclear injuries: (**i**) Dynein-motors are uniformly distributed on the nuclear envelope (NE) and microtubules connecting nucleoskeleton and MTOC in steady-state cells. This organization is not perturbed in presence of small Syn-filaments. (**ii**) Large perinuclear LBs destabilize the microtubules connecting the nucleoskeleton at the immediate locale. Simultaneously, dynein is misrecruited onto the large perinuclear LBs. Partial co-localization of the LBs and the nuclear lamina is also observed. (**iii**) The dynein force working along the distorted microtubules do not anymore connect the nucleoskeleton and creates a misbalanced cytoskeleton tension on the rest of the nucleoskeleton. (**iv**) This results in NE-squeezing that tears the nuclear lamina. (**v-vi**) Pharmacological inhibition of dynein motor or preventing dynein misrecruitment on LBs by knocking down BICD2 relaxes the deregulated tension on NE and re-establishes nucleoskeleton symmetry. **9B. Model depicting two distinct α-Synuclein inclusion bodies in the same cell offering unique homeostatic consequences:** (**i**) Syn-aggresomes positioned on MTOC form spontaneously and assist UPS mediated degradation of the protein. (**ii-iii**) Syn-filaments can be induced by the addition of seeds in the same neurons or mitotic cells. Large perinuclear LBs, but not the small peripheral, Syn-filaments are recognized by PQC factors. Destabilizing Syn-aggresomes by disrupting microtubules accelerates Syn-filament biogenesis. (**iv**) Syn-filaments remain UPS-degradable in aggresome mediated manner upon withdrawal of seeds or after stopping α-Synuclein expression. (**v**) In case of continued polymerization, repurposing of PQC-factors to large LBs oversaturates proteostasis resulting in aggregation of other misfolding-prone proteins; in the current example FlucDM-mCherry (red). Also, large LBs associate with nuclear envelope, weaken and tear the nuclear lamina. Saturated PQC and deregulated transcription in the damaged nuclei in combination increases cellular vulnerability to proteostasis stress.

## Discussion

The name ‘Synuclein’ was derived from the first detection of the protein at the synaptic vesicles (‘Syn’) and its association with nuclear envelope (‘nuclein’) but the association of soluble α-Synuclein with nuclear envelope was considered to be an antibody-staining artefact later (64, 65). Since then, many beneficial or deleterious outcomes of nuclear translocation of α-Synuclein have been described but its association with NE was not confirmed (66–69). Justifying the nomenclature, our study confirms that the LBs associate with and distort nuclear lamina due to an unbalanced mechanical tension between cytoskeleton and nucleoskeleton. This deregulates transcription of stress chaperones and renders the cells vulnerable to proteostasis stresses (**Fig. 9B**). Recently, lamina disintegrities have been reported in Tau and mHtt aggregation models [28, 41, 42]. GWAS studies indicated that a genetic polymorphism in LmnB1 is associated with poor cognitive outcomes in Parkinson’s Disease (70). We find extensive lamina disintegrities in post-mortem PD-brain sections. Together, we propose that amyloid diseases including the LB-related α-Synuclein-proteinopathies may be considered as a sub-group of Laminopathies.

## A cell culture model for the biochemistry of LB-like inclusions

Polymerization of recombinant α-Synuclein into amyloid-filaments is a template dependent self-sufficient process *in vitro* (29). Our results in primary neurons and Hek293T cells suggest that polymerization of α-Synuclein into LB-like filamentous IBs remains a protein-intrinsic process *in cellulo*, does not depend on the endogenous or exogenous source of the protein, can propagate through mitotic divisions, and not restricted to any cell-type specific infrastructure for perinuclear localization. Both the cell types recognize the large perinuclear LB-like IBs as protein-waste for disposal and mark with PQC-factors. Recruitment of dynein onto the LB-like inclusions, overlap of large LB-like IBs with nuclear lamina, asymmetric nucleoskeleton, NE-disintegrities, and deregulation of stress response pathways are also consistent in both cells. Thus, the mechanism of IB-biogenesis and the cell’s reaction to the Syn-IBs are remarkably similar between mitotic cells and post-mitotic neurons validating our cell culture model as a convenient ’test-tube’ for robust cellular biochemistry of LB-like inclusions.

## Coexisting Syn-IBs: an attempt to limit LB-overgrowth, kinetic failure, and associated homeostatic costs

Our results suggest that the homeostatic management of α-Synuclein is co-ordinated by two co-existing IBs in mouse primary neurons and mitotic cells. LB-biogenesis is not required for PQC-management of α-Synuclein. Rather, Syn-aggresomes are sufficient to balance α-Synuclein protein level in cells. LB-biogenesis is initiated in aging neurons only if endogenous α-Synuclein undergoes liquid–liquid phase separation (LLPS) to nucleate amyloidogenesis (26, 71). This may happen due to increased neuronal abundance of α-Synuclein because of gene-duplication or due to mutations in α-Synuclein that accelerates LLPS of the protein (26, 72). Otherwise, LB-biogenesis may be triggered due to propagation of amyloid seeds from one cell to another (73). Our results suggest that turnover of α-Synuclein in Syn-aggresomes serves as the rate limiting mechanism for LB-biogenesis. The kinetics of LB-biogenesis outpaces the turnover only if amyloid-seeding sets in. Although LB-biogenesis is not spontaneously intended for quality control, efficient self-assembly of smaller Syn-filaments into larger LBs avoids co-insolubility of other cellular proteins. Further, LBs may not be the self-sufficient degradation sites but allow conditional disintegration and degradation of α-Synuclein amyloids through the co-existing Syn-aggresomes. Thus, LBs self-quarantine like passive amyloid IPODs but remain UPS-degradable, and we term LBs according to these unorthodox PQC-qualities as SInQCs – <u>S</u>eeding-based <u>In</u>clusions for <u>Q</u>uality <u>C</u>ontrol (**Fig. 9B and S9**).

We find that proteostasis collapse is a late event in LB-containing cells correlated with LB-overgrowth at the perinucleus and nuclear lamina injuries. Mitotic cells can repair these injuries are *via* NE-remodelling after division and overcome the homeostatic challenges. In contrast, LB-related NE-injuries are irreparable in post-mitotic neurons. DNA-damage and epigenetic deregulations in neurons are common pathologic manifestations of Parkinson’s Disease (74, 75). Intriguingly, it is recently reported that post-mitotic neurons in transgenic *Drosophila* activate cell cycle when LmnB1 is depleted at the NE (76). Medium spiny neurons in transgenic mice with mHtt IBs also co-exhibit NE-injuries and cell cycle markers but trigger neurodegeneration via apoptotic pathways (77–79). These may indicate failed attempts to repair NE-injuries via NE-remodelling by triggering division in post-mitotic cells. We posit that controlled activation of ectopic mitosis may repair the injured NE in LB-containing neurons as long as cell death pathways can be controlled.

## Methods

### Expression Constructs

*SNCA* was PCR-amplified from pRK172/α-synuclein (80) and cloned into pcDNA4/TO-EGFP using Kpn I and Xho I (Thermo Scientific). *SNCA* variants (A30P, A53T and A30P+A53T (DM)) were prepared by site directed mutagenesis using primers in **Methods Table S1**. To generate FlucDM construct, FlucDM was PCR amplified from pCI-neo FlucDM EGFP (81) and cloned into pcDNA3.1-mCherry (82) by using Kpn I and Xba I. To generate SNCA(DM)-EGFP-NLS construct, NLS sequence was amplified from pcDNA3.1 Syn^NLS^ obtained from Dr. Mel B. Feany as a gift (67) and cloned into pcDNA4/TO SNCA(DM)-EGFP using BsrG1 and Xba1 (NEB). To prepare LmnB1 over-expressing construct, *lmnb1* gene was PCR amplified from cDNA prepared from Hek293T cells and cloned into pCDNA3.1-mCherry using Not I and Apa I (NEB). pAAV-CAG-H2B tdTomato was obtained from Addgene (Plasmid#116870, (83)). For protein purification, *SNCA* and its mutants (A30P, A53T and DM) were cloned into pET-28a(+) expression vector using the restriction enzymes Hind III and Xho I.

### Protein Purification

SNCA-pET-28a(+) constructs were transformed into *E. coli* BL21(DE3) and induced by adding 1 mM IPTG (Sigma) at 0.6 OD_600_. Cells were pelleted 3 hr after induction, resuspended in TNE buffer (50 mM Tris, 150 mM NaCl, 10 mM EDTA pH 8), and frozen at– 80°C until use. Frozen cell pellets were incubated at 95°C for 10 min and centrifuged at 18,000 g. The supernatant was subjected to streptomycin and ammonium sulphate precipitation. The precipitate was dissolved in HEPES buffer (50 mM HEPES, 100 mM KCl pH 7) and size exclusion chromatography was performed using HiLoad 16/600 Superdex 75 pg Column (GE28-9893-33, GE Healthcare) attached to BioLogic DuoFlow Chromatography system (Bio-Rad). After purification, α-Synuclein and mutants were concentrated using 10 kDa Amicon (Merck Millipore) up to 1-4 mg/ml in HEPES buffer as per requirement.

### Preformed fibril (PFF) generation

1 ml of concentrated protein (1mg/ml) was aliquoted into 1.5 ml tubes (LoBind, Eppendorf) and kept on Eppendorf ThermoMixer C at 37°C with 1500 rpm for 48 hr. Fibril formation was checked using a small fraction in ThT assay and the formed fibrils were sonicated using 3 mm thick probe sonicator (cat: VCX 130 PB, SONICS Vibra-Cell) for 1 min with setting: 5 second ON, 2 second OFF, 15 % amplitude. Sonicated PFF were then aliquoted into 50 and 100 µl volume and stored in – 80°C until use. The PFF stored at – 80°C were stable for 5-6 months with similar aggregation propensity in cell culture.

### Quantitative RT-PCR

Total RNA was prepared using NucleoSpin RNA kit (cat: 740955, Macherey-Nagel) as per manufacturer’s protocol. RNA concentration was measured using Nanodrop 2000 spectrophotometer (Thermo Scientific). For cDNA synthesis, 2 μg of total RNA was used along with SuperScript III Reverse Transcriptase (cat:18080093, Invitrogen) in a final volume of 20 μl. Quantitative PCR was carried out in Applied Biosystems 7900 HT Fast Real-Time PCR System with iTaq^®^ Universal SYBR^®^ Green Supermix (cat:172-5124, Bio Rad) using RT-PCR primers (**Methods Table S2**).

### Cell Culture, transfection and preparation for microscopy

Hek293T cells were maintained in Dulbecco’s Modified Eagle’s Medium (Gibco) supplemented with 10 % fetal bovine serum (FBS) (Gibco) and 90 U/ml penicillin (Sigma) 50 µg/ml streptomycin (Sigma) at 37°C and 5 % CO_2_. Transfection of cells was performed with Lipofectamine 3000 reagent (Invitrogen) as per manufacturer’s protocol. For stable cell line preparation, pcDNA6/TR and pcDNA4/TO system was used (Life Technologies). 24 hr after transfection with pcDNA6/TR, 10,000 cells were seeded in a 100 mm dish and selected with 5 µg/ml Blasticidin S HCl (Invitrogen). Colonies formed after 2-3 weeks were analyzed for anti-TET repressor by western blots (data not shown). Positive clone for pcDNA6/TR was transfected with various pcDNA4/TO constructs and selected with 200 µg/ml Zeocin (Invitrogen). Surviving colonies after 2-3 weeks were tested for induction by adding 1 µg/ml Doxycycline (MP Bioscience) by fluorescence microscopy and western blot. Positive clones were frozen and/or maintained for further experiment.

For LB-like IBs formation, wild type SNCA-EGFP (and other variants) stable cell lines were induced by 1 µg/ml Doxycycline for 24 hr. Next day, induced cells were trypsinized and subcultured into various dishes according to **Methods Table S3** for 6 days. 1 µM PFF and 1 µg/ml doxycycline were added after 6 hr of cell plating.

For SILAC based mass spectrometry – cells were grown in SILAC DMEM (Thermo Scientific) supplemented with 10% dialyzed FBS (Gibco), 90 U/ml penicillin (Sigma), 50 µg/ml streptomycin (Sigma) and either Light [L-Lysine 2HCl / L-Arginine HCl (Lys0/Arg0)] or Medium [L-Lysine 2HCl (4,4,5,5-D4) / L-Arginine HCl (^13^C_6_) (Lys4/Arg6)] or Heavy [L-Lysine 2HCl (^13^C_6_, ^15^N_2_) / L-Arginine HCl (^13^C_6_, ^15^N_4_) (Lys8/Arg10)] isotopes of lysine and arginine (Thermo Scientific).

For Live Cell Microscopy – Cells were grown on Lab-Tek 2-well chambered cover glass (Nunc) under normal growth condition and imaged on OLYMPUS FV3000 Laser scanning microscope at 37°C and 10 % CO_2_. For Live Cell Staining – MitoTracker™ Red CMXRos (Invitrogen) staining was done at a final concentration of 0.5 µM in culture media by incubating cells for 30 min at normal growth conditions. Slides were prepared after acetone/methanol fixation by counterstaining with DAPI (Sigma). For ER and Lysosome staining, cells grown on 2-well chambered cover glass (Nunc) were treated with 100 nM ER-Tracker™ Blue-White DPX (Invitrogen) and 1µM LysoTracker™ Red DND-99 (Invitrogen), respectively, in culture media for 30 min and taken for imaging without fixation using OLYMPUS FV3000 microscope.

For siRNA Knockdown-After 72 hours of PFF incubation in cells grown on cover glass, cells were transfected with 40 nM scrambled (Invitrogen Silencer™ select negative control, cat # 4390843) or 50 nM BICD2 siRNA (Invitrogen Silencer™ select BICD2, Assay ID-s23497, cat# 4392420) using Lipofectamine® RNAiMAX (Invitrogen, Cat#13778-150) and continued incubation for next 72 hours before fixing.

Acetone/Methanol fixation – Cells grown on cover glass were washed with PBS to remove medium. Pre-chilled acetone and methanol in ratio 1:1 was added to the cells and plate was incubated at -20 °C for 5 min followed by PBS wash.

Paraformaldehyde fixation –Cells were incubated with 4% paraformaldehyde solution at room temperature for 15 min followed by PBS wash. 0.1% Triton X100 in PBS was used for cell permeabilization prior to immunostaining.

DiIC_16_ staining-For lipid staining, 8µM of DiIC_16_ added to the fixed cells post permeabilization and incubated for 15 min of ice followed by PBS wash (84).

### Primary Neuron Culture

Hippocampal Primary neurons were isolated from E17-E19 C57BL/6 mouse brains (Charles River Laboratories) as described in Seibenhener *et al* (85). Approx. 90000 cells were plated on poly-D-lysine coated 18 mm glass coverslips. Neurobasal Feeding Media was replaced every third day. For Syn-filament generation, 100 nM PFF was added at 5 days *in vitro* (DIV) and incubated for 10 days at described in Volpicelli-Daley *et al* (17). All experiments were performed as per the standard protocols and procedures approved by the Institutional Animal Ethics Committee (IAEC). Primary neurons were fixed on 15 DIV with 1:1 acetone and methanol chilled at -20 °C and immune stained with appropriate antibodies.

### Immunofluorescence microscopy

Cells were grown on 18 mm glass coverslips in 12 well plate (Nunc) and fixed as explained above. After fixation, the coverslips were incubated with 5% BSA prepared in PBS with 0.05% Tween-20 (PBST) for 30 min at RT with constant shaking. Primary antibodies (**Methods Table S4**) were added and coverslips were incubated at 4°C overnight followed by PBST wash. Next day, secondary antibodies were added to the coverslip and incubated for 1 hr at room temperature. Antibody dilutions were prepared in 0.1% BSA with PBST as mentioned in **Methods Table S4** Coverslips were mounted on slides using 10 µl Antifade Mounting Medium (cat: H-1000, Vectashield) after DAPI staining. For Actin filament staining, Fixed cells were stained with 1:1000 dilution of ActinRed™ 555 ReadyProbes™ Reagent (Rhodamine phalloidin) (cat: R37112, Thermo scientific) for 30 min at RT. Imaging was done in either Leica TCS SP8 or ZEISS LSM 880 laser scanning microscope.

Pearson coefficients for the colocalization analysis are presented in **Supplementary Note Colocalization Figures and Tables**.

### Immunofluorescence (Paraffin-Embedded Sections)

The human post-mortem thalamus sections of a Parkinson’s Disease (PD) patient (BioChain Institute Inc., USA Cat# T2236079Par) and a control (BioChain Institute Inc., USA Cat#T2234079) were treated with Xylene (Himedia Cat#AS080-2.5l) twice for 10 min. Followed by washes with ethanol (100%, 95%, 70 %) and H2O for 10 min each. Sections were permeabilization with 0.2% Triton X-100 in PBS for 5 min, followed by antigen retrieval in Citrate buffer (10mM Trisodium citrate, pH 6, 0.05 % Tween20). After blocking with 4% horse serum (Gibco) in PBST, sections were incubated overnight with primary antibodies at 4°C followed by incubation with suitable secondary antibody for 1hr at RT. The antibody dilutions were prepared in 4% horse serum in PBST. Sections were then mounted using ProLong Diamond antifade reagent with DAPI (Thermo Fisher Scientific).

### Cell Viability and Cytotoxicity assay

MTT (Sigma) solution (0.5 mg/mL in growth medium) was added to cell culture plates followed by incubation for 3 hr under normal growth conditions. Media was removed and formazan crystals were dissolved in DMSO. Absorbance was measured at 570 nm using Multiskan GO Plate Reader (Thermo Scientific). Cell toxicity was measured using Pierce LDH Cytotoxicity Assay Kit (Thermo Scientific) as per manufacturer’s protocol. Equal volume of culture media and reaction mix was taken in a fresh plate and incubated for 30 min in dark at RT. Absorbance was recorded at 490 nm in Multiskan GO Plate Reader (Thermo Scientific).

### Cell lysis, SDS-PAGE and Western Blotting

Cell pellets were lysed in TBS-T lysis buffer (1% Triton X-100 in 50 mM Tris-HCl, 150 mM KCl, and a cocktail of protease inhibitors (Roche)) at 4°C and incubated for 1 hr with intermittent vortexing. Lysed cells were centrifuged at 12,000 g for 15 min at 4°C, and the supernatant was collected as soluble fraction. The pellet was washed twice with 1×PBS and boiled in 4×SDS loading buffer (0.2 M Tris–HCl pH 6.8, 8 % SDS, 0.05 M EDTA, 4 % 2-mercaptoethanol, 40 % glycerol, 0.8 % bromophenol blue) for 15 min to obtain the insoluble fraction. Total fraction was prepared by directly boiling the cell pellet in 4×SDS loading buffer for 15 min. For non-nuclear, nuclear, and nuclear insoluble fraction preparation, cell pellets were processed using NE-PER Nuclear and Cytoplasmic Extraction Kit (cat:78833, Thermo Scientific). Protein estimation was carried out by Amido Black Protein assay. Protein fractions were separated by SDS-PAGE and transferred onto 0.2 μm PVDF membrane (Bio-Rad) for 90 min at 300 mA using the Mini-Trans Blot cell system (Bio-Rad). Membranes were probed by primary and secondary antibodies (**Methods Table S4**) and imaged using documentation system (Chemi-Smart 5000, Vilber Lourmat).

### Sample Preparation for Mass Spectrometry

**Hek293T** - For total, soluble and insoluble proteome, cell-extracts were prepared as per the experimental scheme in **Fig. S3G**. Cells were pelleted down at day 3 and 6. Equal number (∼1 million) of L-, M-, and H-labeled cells were pooled and lysed together. Proteome fractions were separated on NuPAGE 4%–12% Bis–Tris Protein Gels (Invitrogen) in MES buffer (100 mM MES, 100 mM Tris–HCl, 2 mM EDTA, 7 mM SDS) at 200 V for 40 min, fixed and stained with Coomassie brilliant blue. Preparation of gel slices, reduction, alkylation, and in-gel protein digestion was carried out as described by Shevchenko et al.

(86). Finally, peptides were desalted and enriched according to Rappsilber et al. (87) and stored in -20°C until mass spectrometry analysis. Similarly, equal number (∼ 1.5 million) of L-and H-labeled cells were pooled together to prepare non-nuclear, nuclear and nuclear insoluble fraction as explained in experimental scheme in **Fig. S5E**.

**Primary Neurons** – Primary neurons were directly dissolved in 4% SDS lysis buffer, Neurons were heat stressed as explained in experimental scheme (**Fig. 4J**). The lysate were precipitated by overnight incubating with chilled acetone at -20°C. The protein precipitate was then resuspended in 8M Urea with 7.5 mM DTT in 50 mM ammonium bicarbonate by vortexing for 10 min at RT. Resuspended proteins were reduced incubating with 45 mM DTT for 30 min at RT, followed by alkylation with 10 mM Iodoacetamide for 30 min in dark at RT. For trypsin digestion, Urea in the sample was diluted 10-fold and trypsin was added 1:20 ratio (15ng/µl in 25mM ammonium bi-carbonate/1mM calcium chloride) for 16 hours at 37°C. After 16 hours, Trifluoroacetic acid (10%) was added to stop the trypsin activity and samples were vacuum dried. Dried peptides were dissolved in 5% Acetonitrile/0.1% Trifluoroacetic acid. The peptides were desalted and fractionated using 200 µl Pierce C18 column tips (Merck Millipore).

Peptides eluted from desalting tips were dissolved in 2% formic acid and sonicated for 5 min. Peptides were then analyzed on Q Exactive HF (Thermo Scientific) mass spectrometer interfaced with nano-flow LC system (EASY-nLC 1200, Thermo Scientific). EasySpray Nano Column PepMap^TM^ RSLC C18 (Thermo Fisher) (75 μm× 15 cm; 3 μm; 100 Å) using 60 min gradient of mobile phase [5% ACN containing 0.1% formic acid (buffer A) and 90% ACN (acetonitrile) containing 0.1% formic acid (buffer B)] at flow rate 300 nL/min was used for separation of peptides. Full scan of MS spectra (from m/z 400 to 1750) were acquired followed by MS/MS scans of top 10 peptide with charge state 2 of higher.

The MS-based proteomics data of all these experiments have been deposited to the ProteomeXchange Consortium via the PRIDE partner repository (88) with the dataset identifier PXD028941.

### Peptide Statistics

For peptide identification, raw MS data files were loaded onto MaxQuant proteomics computational platform (Ver. 1.6.10.43) (89) and searched against Swissprot database of Homo sapiens (release 2019 with 20,371 entries) and a database of known contaminants. A decoy version of the specified database was used to adjust the false discovery rates for proteins and peptides below 1%. The search parameters included constant modification of cysteine by carbamidomethylation, enzyme specificity trypsin, and multiplicity set to 3 with Lys4 and Arg6 as medium label and Lys8 and Arg10 as heavy label. Other parameters included minimum peptide for identification 2, minimum ratio count 2, re-quantify option selected, and match between runs with 2-min time window.

SILAC labelled cells downstream analysis and statistics was done by normalizing ratios (H/L and M/L) from two biological repeat experiments was done using Perseus (Ver. 1.6.0.4, 1.6.10.43) (90). Individual experiment H/L ratios for each fraction were converted into log2 space and one-sample t-test was performed to calculate the statistical significance with Perseus. For Z-score calculation, average ratio H/L of 2 repeats was converted into log2 space and normalized using mean and standard deviations for each data set in Perseus.

For Label free quantification, raw spectra files were loaded onto MaxQuant proteomics computational platform (Ver. 1.6.10.43) and searched against SwissProt *Mus Musculus* Fasta database (version November 2022). The search parameters included static and dynamic modification of cysteine by carbamidomethylation and dynamic modification of methionine by oxidation, respectively, enzyme specificity of trypsin allowing for up to 2 missed cleavages. Precursor mass tolerance was set to 5 ppm and fragment mass tolerance

0.02 Da was used. Other parameters included minimum 1 peptide for identification with 6 amino acid length. Percolator (q-value) was used for validating peptide spectrum matches and peptides, accepting only the top-scoring hit for each spectrum, satisfying the cut off values for FDR 1% and Label Free Quantification (LFQ) option was selected.

Log2 fold change was calculated from the ratios of LFQ intensities of Primary Neurons (DIV 15), - PFF (Heat Shock/ Non-Heat Shock) and + PFF (Heat Shock/ Non-Heat Shock) from average LFQ intensities of 3 independent experiments. Heatmaps were prepared using Morpheus (https://software.broadinstitute.org/morpheus) and other graphs were prepared in Perseus, GraphPad prism ver. 9.4.1 or OriginPro 2021b (OriginLab).

**Methods Table S1.**
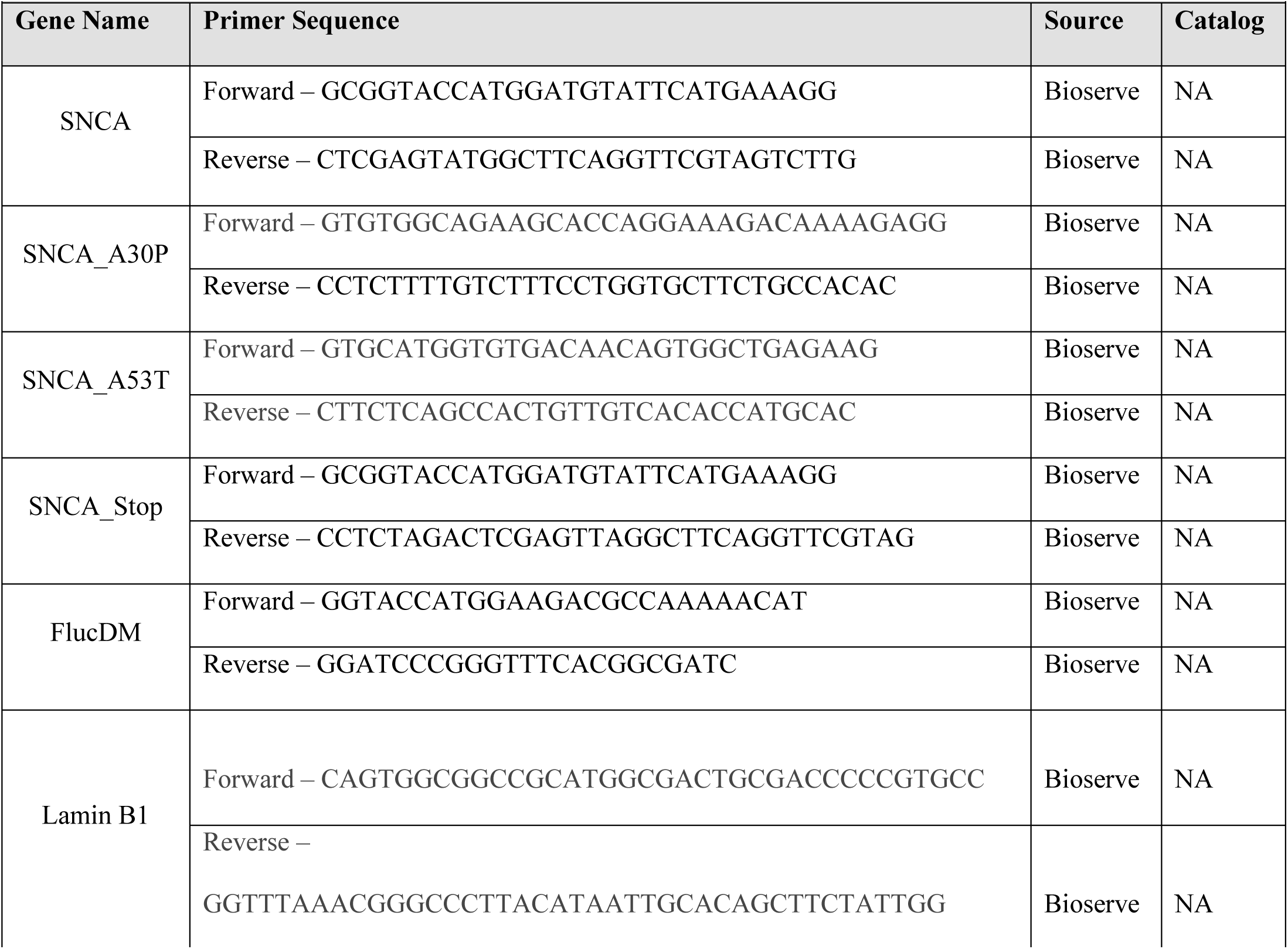
Cloning Primers

**Methods Table S2.**
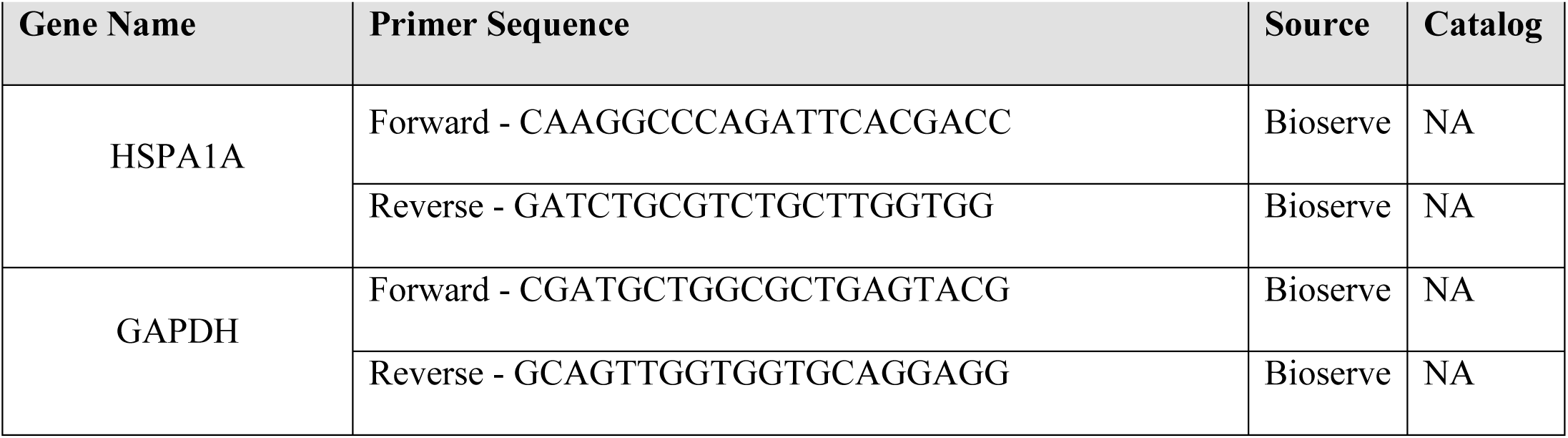
Quantitative RT-PCR Primers

**Methods Table S3.** Cell Number

| Methods Table S3 – Cell Number |  |  |
| --- | --- | --- |
| Format | Media (ml) | Cell number |
| 12-well plate (Nunc) | 1 | 5,000 |
| 6-well plate (Nunc) | 2 | 10,000 |
| 60mm dish (Nunc) | 3 | 15,000 |
| 100mm dish (Nunc) | 10 | 1,00,000 |
| 2-chambered cover glass (Nunc) | 1 | 5,000 |

**Methods Table S4.**
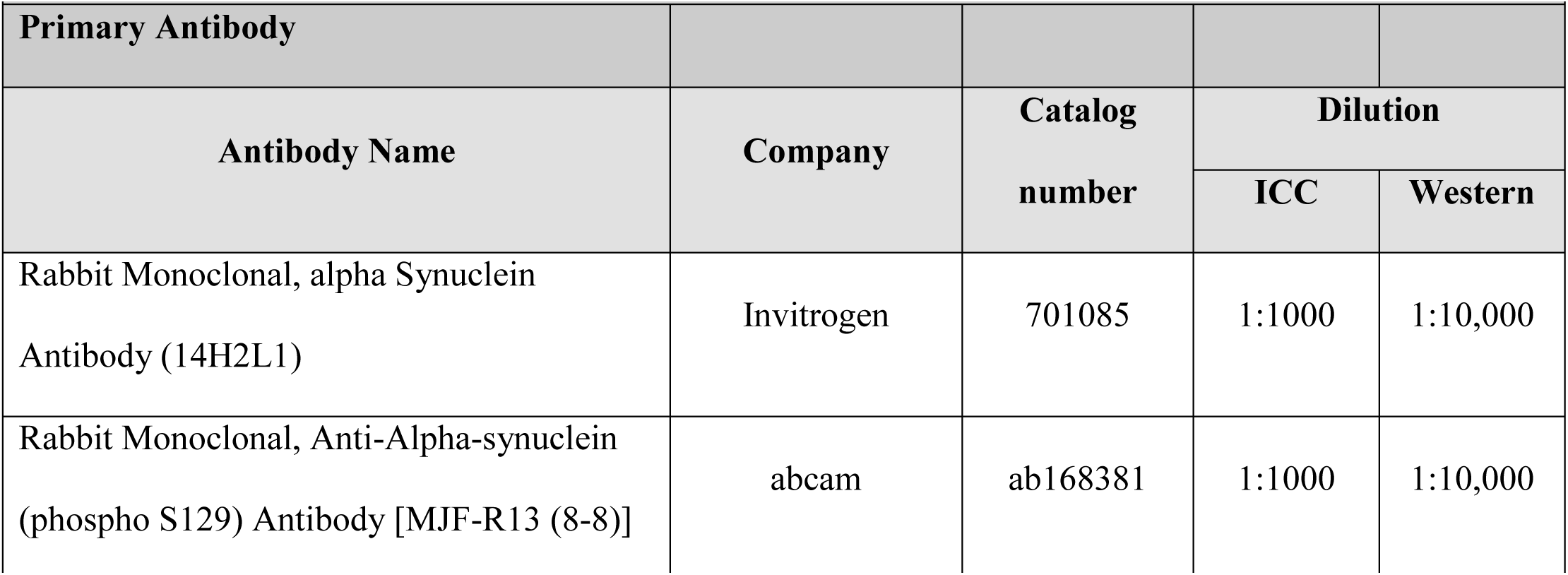
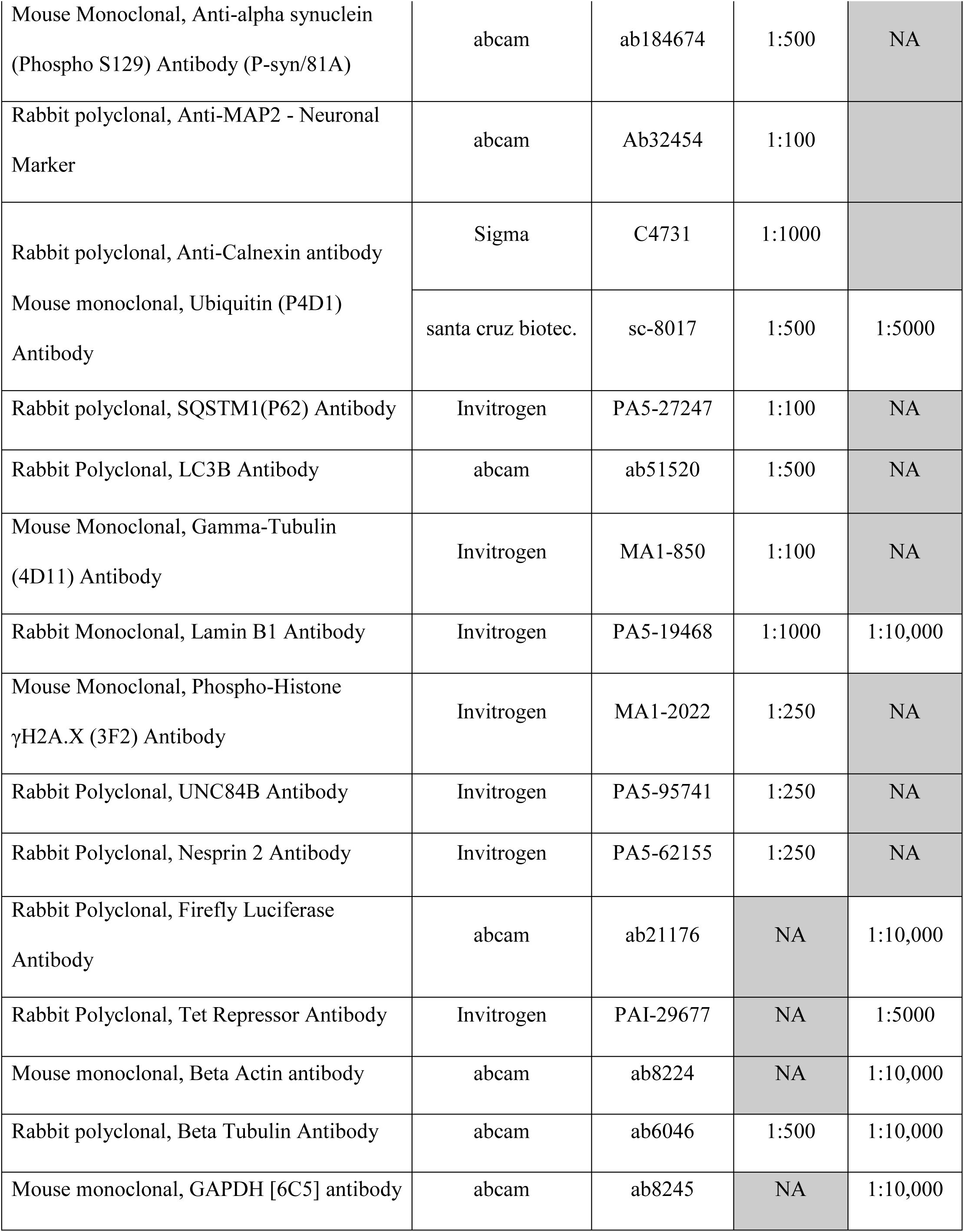
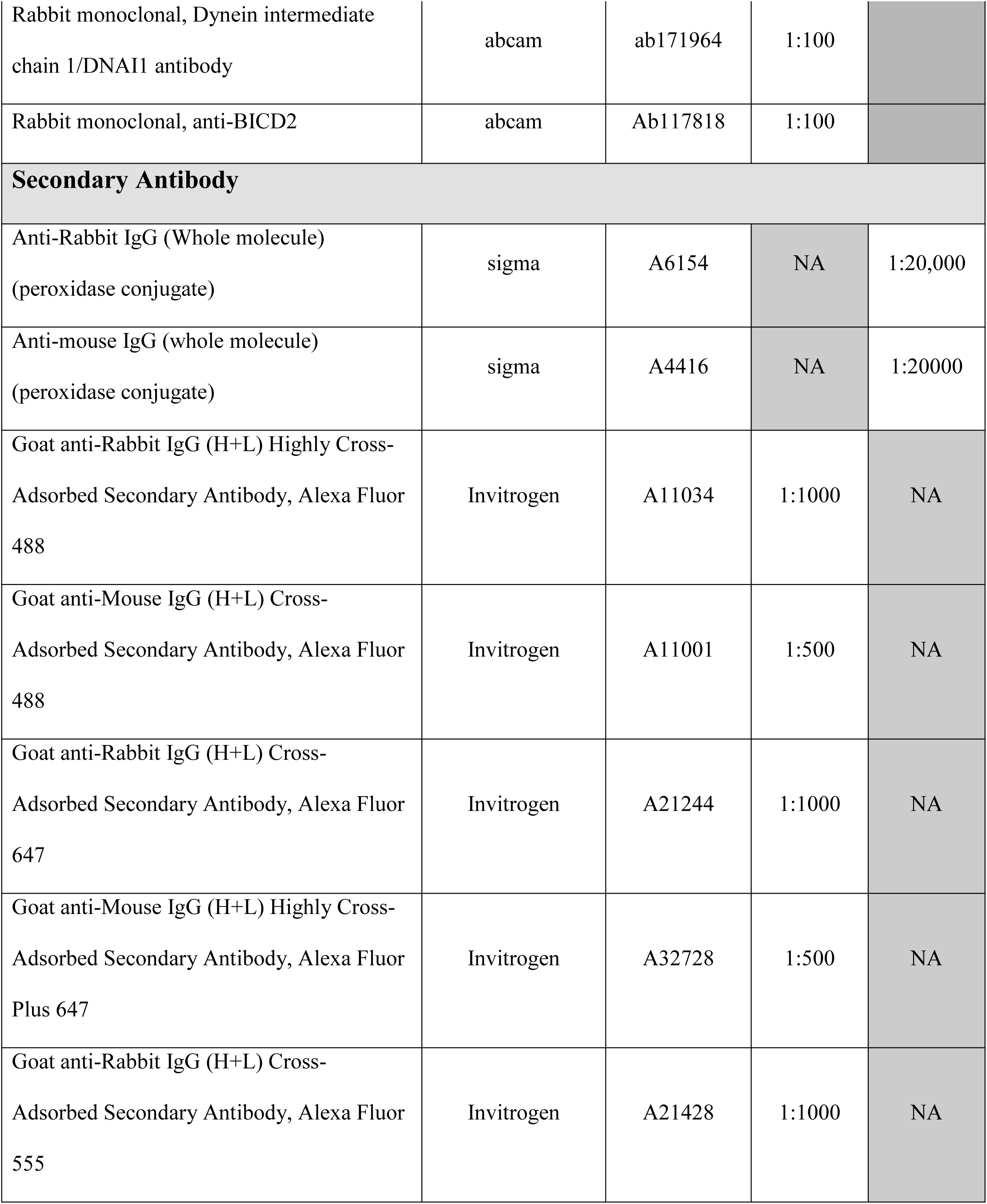
Antibody List

| Methods Table S4 - Antibody List |  |  |  |  |
| --- | --- | --- | --- | --- |
| Primary Antibody |  |  |  |  |
| Antibody Name | Company | Catalog number | Dilution |  |
|  |  |  | ICC | Western |
| Rabbit Monoclonal, alpha Synuclein Antibody (14H2L1) | Invitrogen | 701085 | 1:1000 | 1:10,000 |
| Rabbit Monoclonal, Anti-Alpha-synuclein (phospho S129) Antibody [MJF-R13 (8-8)] | abcam | ab168381 | 1:1000 | 1:10,000 |
| Mouse Monoclonal, Anti-alpha synuclein (Phospho S129) Antibody (P-syn/81A) | abcam | ab184674 | 1:500 | NA |
| Rabbit polyclonal, Anti-MAP2 - Neuronal Marker | abcam | Ab32454 | 1:100 |  |
| Rabbit polyclonal, Anti-Calnexin antibody | Sigma | C4731 | 1:1000 |  |
| Mouse monoclonal, Ubiquitin (P4D1) Antibody | santa cruz biotec. | sc-8017 | 1:500 | 1:5000 |
| Rabbit polyclonal, SQSTM1(P62) Antibody | Invitrogen | PA5-27247 | 1:100 | NA |
| Rabbit Polyclonal, LC3B Antibody | abcam | ab51520 | 1:500 | NA |
| Mouse Monoclonal, Gamma-Tubulin (4D11) Antibody | Invitrogen | MA1-850 | 1:100 | NA |
| Rabbit Monoclonal, Lamin B1 Antibody | Invitrogen | PA5-19468 | 1:1000 | 1:10,000 |
| Mouse Monoclonal, Phospho-Histone γH2A.X (3F2) Antibody | Invitrogen | MA1-2022 | 1:250 | NA |
| Rabbit Polyclonal, UNC84B Antibody | Invitrogen | PA5-95741 | 1:250 | NA |
| Rabbit Polyclonal, Nesprin 2 Antibody | Invitrogen | PA5-62155 | 1:250 | NA |
| Rabbit Polyclonal, Firefly Luciferase Antibody | abcam | ab21176 | NA | 1:10,000 |
| Rabbit Polyclonal, Tet Repressor Antibody | Invitrogen | PAI-29677 | NA | 1:5000 |
| Mouse monoclonal, Beta Actin antibody | abcam | ab8224 | NA | 1:10,000 |
| Rabbit polyclonal, Beta Tubulin Antibody | abcam | ab6046 | 1:500 | 1:10,000 |
| Mouse monoclonal, GAPDH [6C5] antibody | abcam | ab8245 | NA | 1:10,000 |
| Rabbit monoclonal, Dynein intermediate chain 1/DNAI1 antibody | abcam | ab171964 | 1:100 |  |
| Rabbit monoclonal, anti-BICD2 | abcam | Ab117818 | 1:100 |  |
| **Secondary Antibody** |  |  |  |  |
| Anti-Rabbit IgG (Whole molecule)<br>(peroxidase conjugate) | sigma | A6154 | NA | 1:20,000 |
| Anti-mouse IgG (whole molecule)<br>(peroxidase conjugate) | sigma | A4416 | NA | 1:20000 |
| Goat anti-Rabbit IgG (H+L) Highly Cross-Adsorbed Secondary Antibody, Alexa Fluor 488 | Invitrogen | A11034 | 1:1000 | NA |
| Goat anti-Mouse IgG (H+L) Cross-Adsorbed Secondary Antibody, Alexa Fluor 488 | Invitrogen | A11001 | 1:500 | NA |
| Goat anti-Rabbit IgG (H+L) Cross-Adsorbed Secondary Antibody, Alexa Fluor 647 | Invitrogen | A21244 | 1:1000 | NA |
| Goat anti-Mouse IgG (H+L) Highly Cross-Adsorbed Secondary Antibody, Alexa Fluor Plus 647 | Invitrogen | A32728 | 1:500 | NA |
| Goat anti-Rabbit IgG (H+L) Cross-Adsorbed Secondary Antibody, Alexa Fluor 555 | Invitrogen | A21428 | 1:1000 | NA |

**Methods Table S5.** Laser Intensity

| Methods Table S5 - Laser Intensity |  |  |  |  |  |  |
| --- | --- | --- | --- | --- | --- | --- |
|  | Argon (488) (30%) |  | HeNe(633) |  | DPSS(561) |  |
| Figure | Intensity % | Smart Gain | Intensity % | Smart Gain | Intensity % | Smart Gain |
| Fig. 1Ai, Fig. S1Bi | 7.6 | 968 | 30.17 | 998 |  |  |
| Fig. 1Aii, Fig. S1Bii | 28 | 756 | 57 | 987 |  |  |
| Fig. 1Bi, Fig. S1Ci | 8.8 | 907 | 20.49 | 916 |  |  |
| Fig. 1Bii, Fig. S1Cii | 8.8 | 667 | 20.49 | 916 |  |  |
| Fig. S1A | 6.4 | 770 | 35 | 886 |  |  |
| Fig. S1Di | 13.66 | 955 | 7.2 | 840 |  |  |
| Fig. S1Dii | 13.66 | 955 | 8.46 | 560 |  |  |
| Fig. S1Ei | 6.4 | 882 |  |  | 21.77 | 818 |
| Fig. S1Eii | 13.66 | 955 |  |  | 21.77 | 818 |
| Fig. 5I Left | 7.61 | 792 | 30.17 | 785 |  |  |
| Fig. 5I Right | 7.61 | 895 | 30.17 | 785 |  |  |
| Fig. 5J, Fig. S5J | 28.18 | 691 | 45 | 996 |  |  |
| Fig. S5I | 7.61 | 895 | 30.17 | 785 |  |  |
| Fig. 6B Top, Fig. S6B Left | 1 | 550 | 1 | 464 | Olympus fv3000 confocal Microscope |  |
| Fig. 6B Bottom, Fig. S6B Right | 0.98 | 537 | 1 | 535 |  |  |
| Fig. 6C Left, Fig. S6C Left | 28.18 | 848 | 45.81 | 762 |  |  |
| Fig. 6C Right, Fig. S6C Right | 28.18 | 848 | 45 | 762 |  |  |
| Fig. 6Di, Fig. S6D Left | 28.18 | 945 | 57 | 995 |  |  |
| Fig. 6Dii, Fig. S6D Right | 28.18 | 683 | 57 | 674 |  |  |
| Fig. 6Ei, Fig. S6E Left | 37 | 962 | 57.9 | 854 |  |  |
| Fig. 6Eii, Fig. S6E Right | 28.18 | 954 | 57.9 | 837 |  |  |

## Acknowledgment

The authors acknowledge Dr. Arvind Kumar, CSIR-Centre for Cellular and Molecular Biology (CCMB) and Dr Sumona Chakravarty, CSIR-Indian Institute of Chemical Biology (IICT) for help in primary neuron experiments and Mr. Jerald Mahesh Kumar, Mr. N Sai Ram and Mrs. B Jyothi Lakshmi at CSIR-CCMB animal house facility for maintaining and providing animals. pAAV-CAG-H2B tdTomato and pcDNA3.1 Syn^NLS^ constructs were kind gifts from Dr. P Chandra Shekar, CSIR-CCMB and Dr. Mel B. Feany, Harvard Medical School, USA. DiIC_16_ lipid staining dye was obtained from Dr. Amitabha Chattopadhyay, CSIR-CCMB as a gift. The authors also acknowledge CSIR-CCMB Proteomics facility, Advance microscopy facility and Tissue culture facility.

## This PDF file includes

Figs. S1 to S9

## Other Supplementary Materials for this manuscript include the following

Movies S1 to S5

Tables S1 to S6

**Fig S1.**
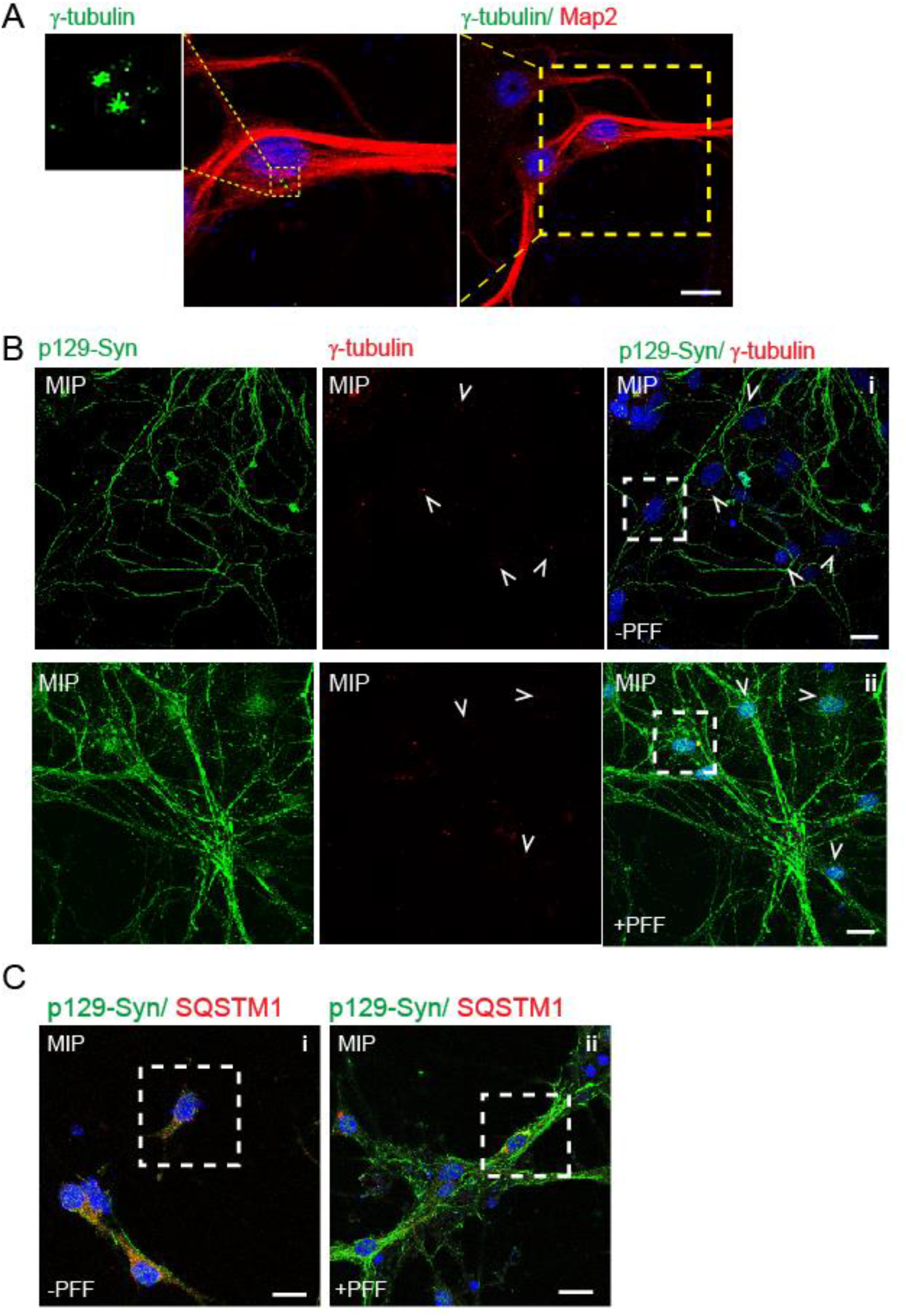
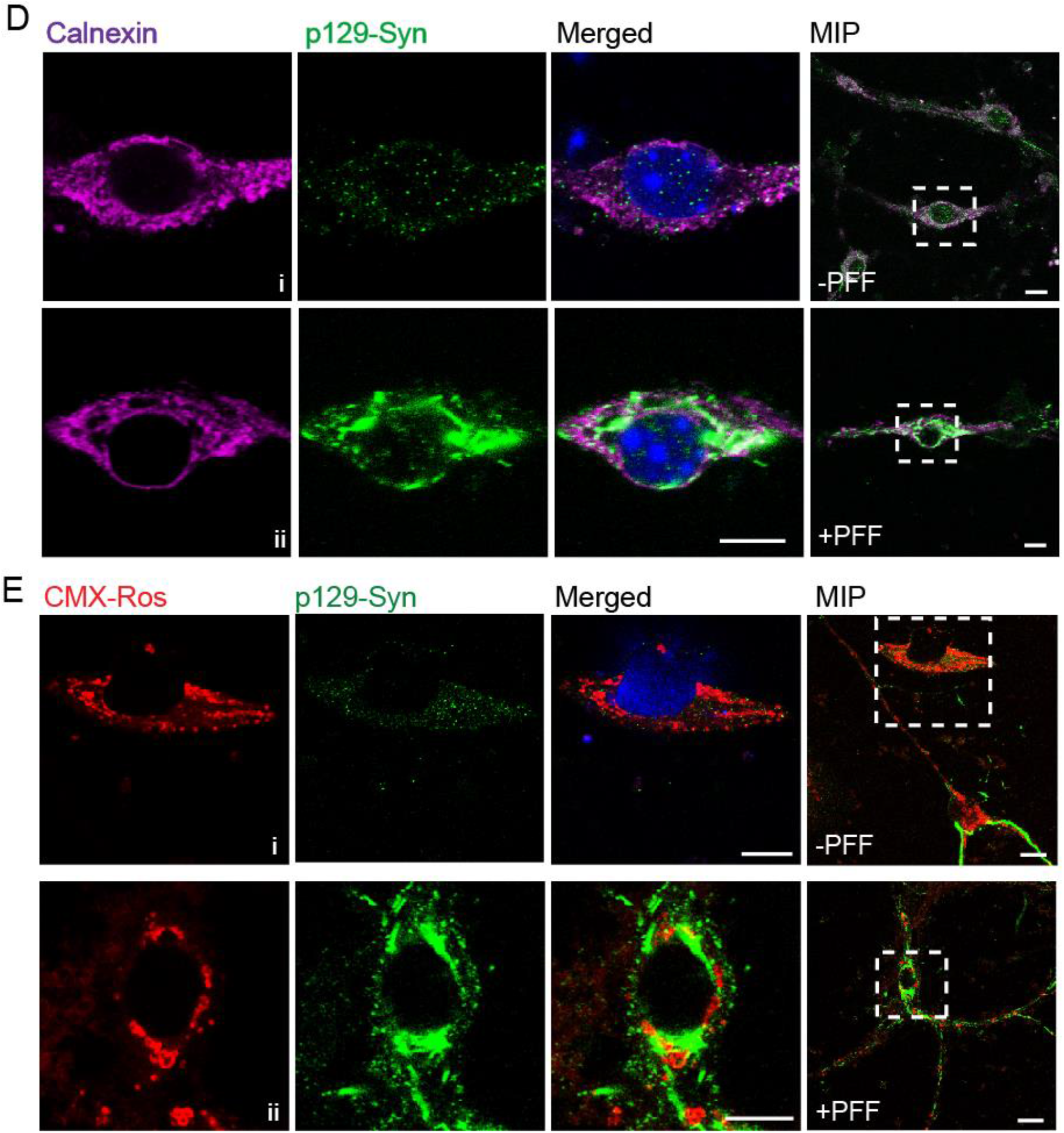
Two unique inclusion bodies of α-Synuclein in primary neurons. **A**: Primary neurons at 15 DIV immunostained with anti-MAP2 and γ-tubulin antibodies. Dotted box indicates zoomed locations. **B-C:** Maximum intensity Projections (MIP) of larger fields for images shown in **Fig. 1A-B. D:** Primary neurons, with and without100 nM PFF from 5 days *in vitro* (DIV) immunostained at 15 DIV with p129-Syn (P-syn/81A) and Calnexin (ER marker) antibody. **E:** Primary neurons with and without PFF treatment at 15 DIV immunostained with p129-Syn (MJF-R13 (8-8)) and stained with CMX-Ros (mitochondria marker). Dotted box indicates zoomed locations. Different channels of single z-plane and Maximum Intensity Projection (MIP) of the larger field is shown. Scale bar-10µm. All images shown in the figure were fixed with Acetone:Methanol.

**Fig. S2:**
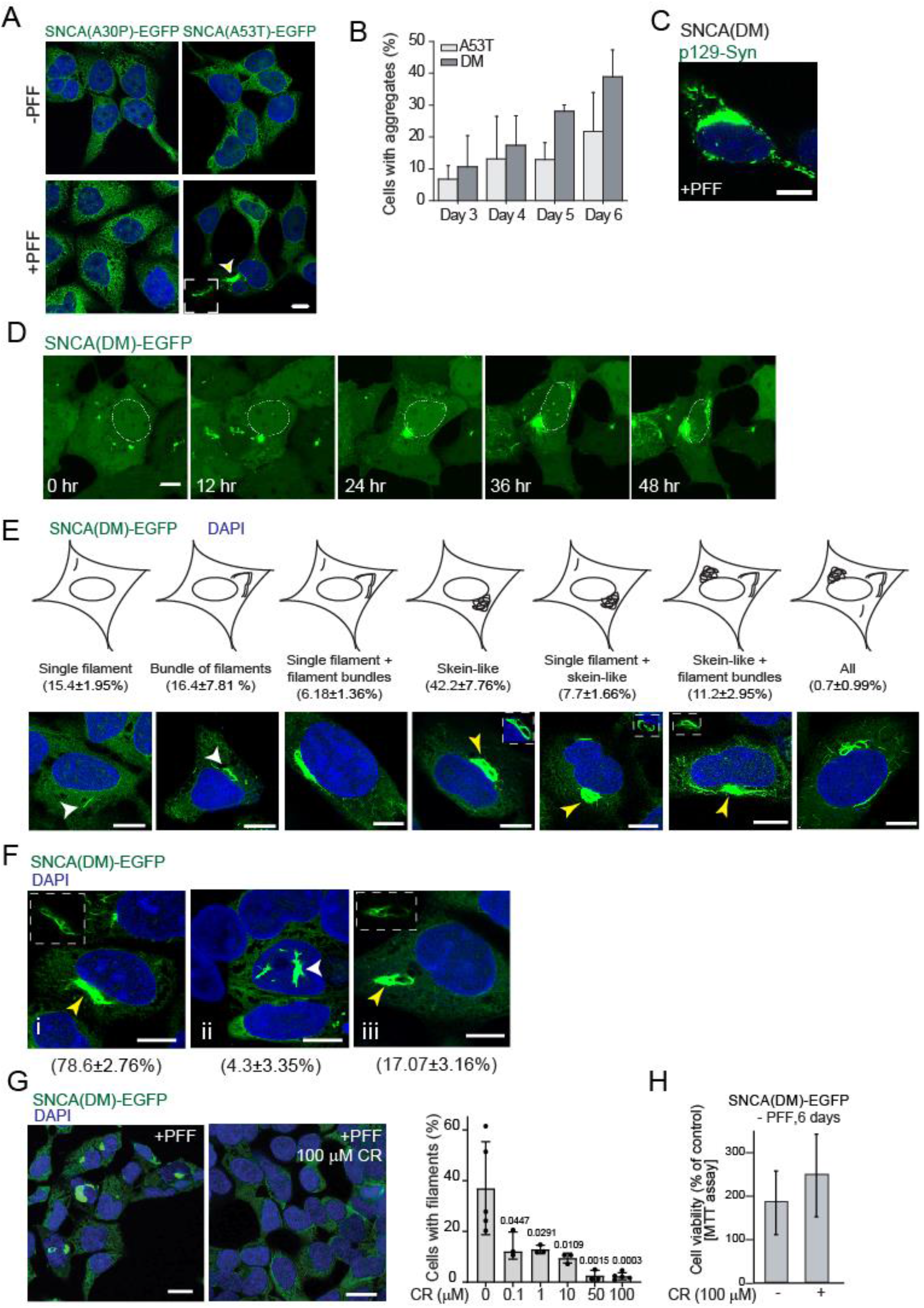
Co-existence of two α-Synuclein IBs is not restricted to post-mitotic neurons. **A**: Hek293T stable lines expressing EGFP-tagged α-Synuclein (SNCA) variants (A30P and A53T) imaged after 6 days of with and without PFF-treatment. **B:** Number of SNCA(A53T)-EGFP and SNCA(DM)-EGFP cells with IBs during the course of PFF-incubation. Approx. 100 cells counted in each experiment. N = 3. Scale bar – 10 µm. **C:** SNCA(DM) Hek293T cells after 6 day PFF treatment stained with p129-SNCA antibody.**D:** Live cell microscopy showing Syn-filaments at different stages of biogenesis. Imaging started at 4^th^ day of the experimental scheme shown in **Fig. 2C**. Dashed circles indicates nuclear boundary. **E**: Morphology of Syn-filaments in 6^th^ day PFF-treated SNCA(DM)-EGFP cells with percent distribution. Approx. 100 cells have been counted in each experiment. N = 3. Insets – low exposure images of the yellow arrow-indicated filamentous IBs, white arrowheads indicate small filaments. **F:** Localization of Syn-filaments in 6^th^ day PFF-treated SNCA(DM)-EGFP cells with percent distribution. Approx. 100 cells have been counted in each experiment. N = 3. Insets – low exposure images of the yellow arrow-indicated IBs. Scale bar – 10 µm**. G: Left:** SNCA(DM)- EGFP cells treated with 1 µM PFF and 100µM Congo Red (CR) for 6 days were fixed and imaged. **Right:** SNCA(DM)-EGFP cells treated with 1 µM PFF and varied concentration of Congo Red (CR) for 6 days. Number of Syn-filament containing cells counted and plotted. Approx. 200 cells were counted in each experiment. Control and 100 µM (N = 5); 0.1 µM, 1 µM, 10 µM and 50 µM (N = 3). Error bar indicates the standard deviation**. H:** Cell viability of 6^th^ day 100 µM Congo Red (CR) treated SNCA(DM)-EGFP cells as estimated by MTT assay. Error bars indicate SD. N=4. **Fig. S2A, C, E F &G** – Acetone:Methanol fixation. **Fig. S2D** – Live cell microscopy without fixation.

**Fig. S3:**
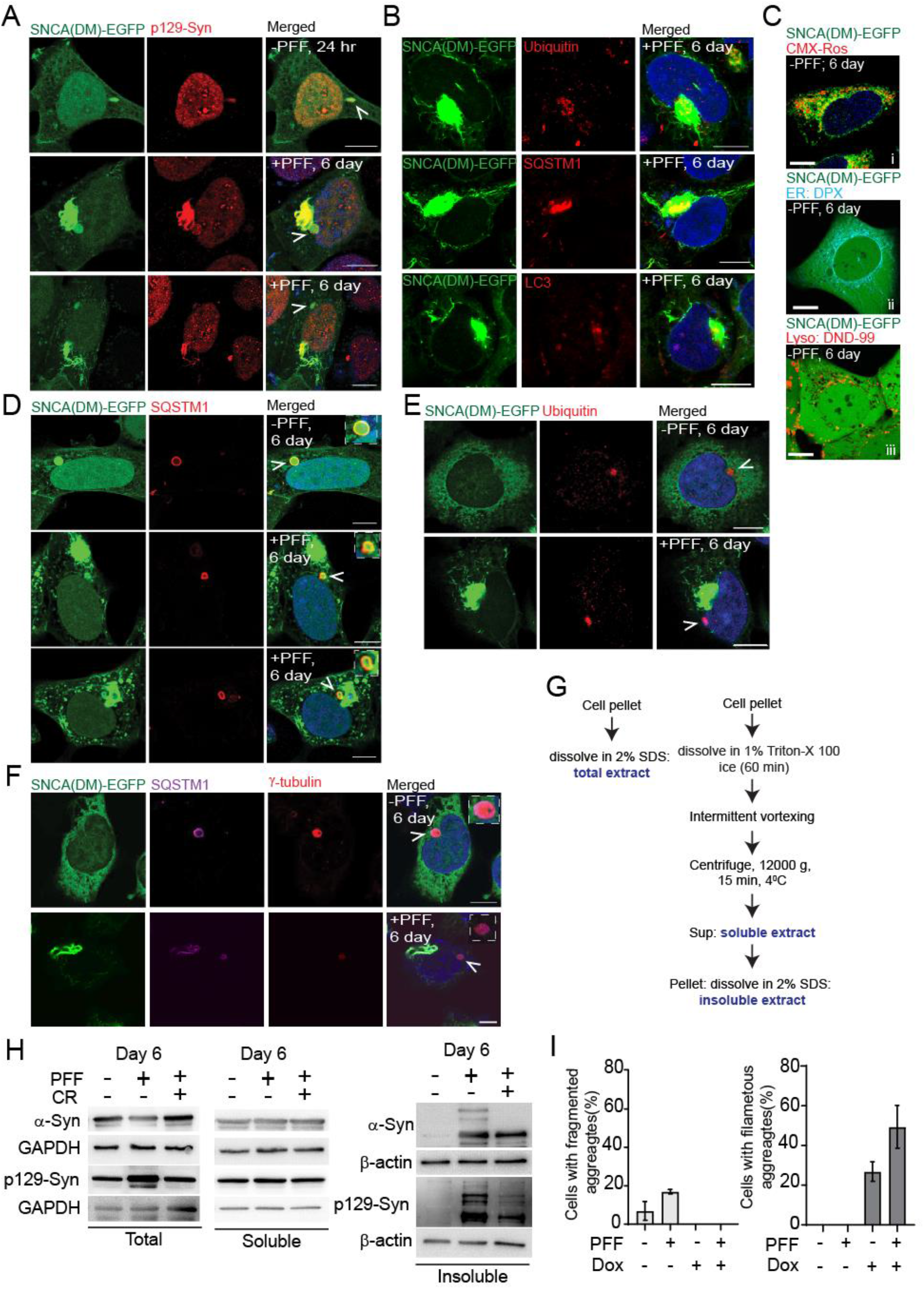
Syn-aggresomes and LB-like IBs offer unique homeostatic opportunities. **A-B**: Microscopic images shown in **Fig. 2E and 2F**– images of fluorescent proteins/ antibodies excited by different wavelengths of lasers shown separately. **C:** 6^th^ day SNCA(DM)-EGFP cells were stained for various cell organelles and imaged. **D-F:** Microscopic images shown in **Fig. 2H, 2I and 2J**– images of fluorescent proteins/ antibodies excited by different wavelengths of lasers shown separately. **G:** Schematic describing the experimental strategies to prepare total, soluble and insoluble cell extracts. **H:** Total, soluble and insoluble fraction prepared from 6^th^ day SNCA(DM)-EGFP cells with Congo Red and probed for α-Syn and p129-Syn. GAPDH and β-actin as loading control. Scale bar – 10 µm. **I:** Number of SNCA(DM)-EGFP cells with fragmented and intact filamentous IBs after 6 days of re-plating upon incubation at indicated conditions. Approx. 100 cells have been counted in each experiment. N = 3. **Fig. S3B, Ci, E-F** – Acetone:Methanol fixation. **Fig. S3A, D** – Paraformaldehyde fixation. **Fig. S3Cii-iii** – Live cell microscopy without fixation.

**Fig. S4:**
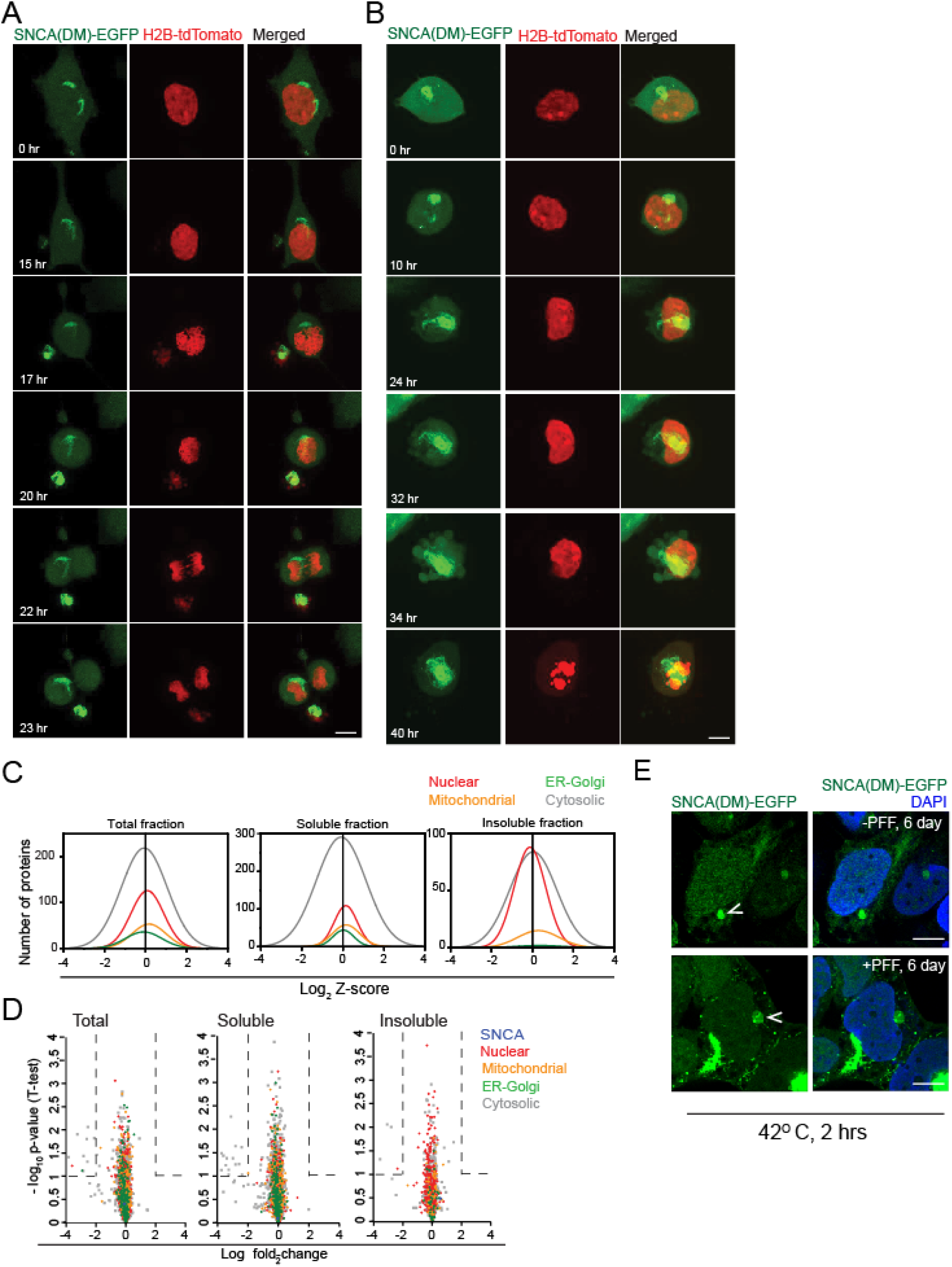
Cellular homeostasis is near saturation in LB-like IB containing cells. **A-B**: Live cell microscopic images shown in Fig. 4B-C – images of fluorescent proteins excited by different wavelengths of lasers are shown separately. Scale bar – 10 µm. **C:** Z-score (SNCA(DM)-EGFP; 6^th^ day, Ratio: +PFF/-PFF) distribution of proteins identified by mass spectrometry according to their subcellular localization (as curated by UniProt) in total, soluble and insoluble fractions. Experimental scheme presented in Fig. S4C**. D:** Volcano plot illustrating proteome equilibrium in 6^th^ day PFF treated SNCA(DM)-EGFP cells. The −log10 (p-value, One-sample *t*-test) is plotted against the log2 (fold change: SNCA(DM)-EGFP; 6^th^ day +PFF/-PFF). **E:** 6^th^ day SNCA(DM)-EGFP cells (-/+) PFF treatment after 2 hr heat shock at 42°C were fixed with paraformaldehyde and imaged. Stick arrows indicate Syn-aggresome. Scale bar-10 µm.

**Fig. S5:**
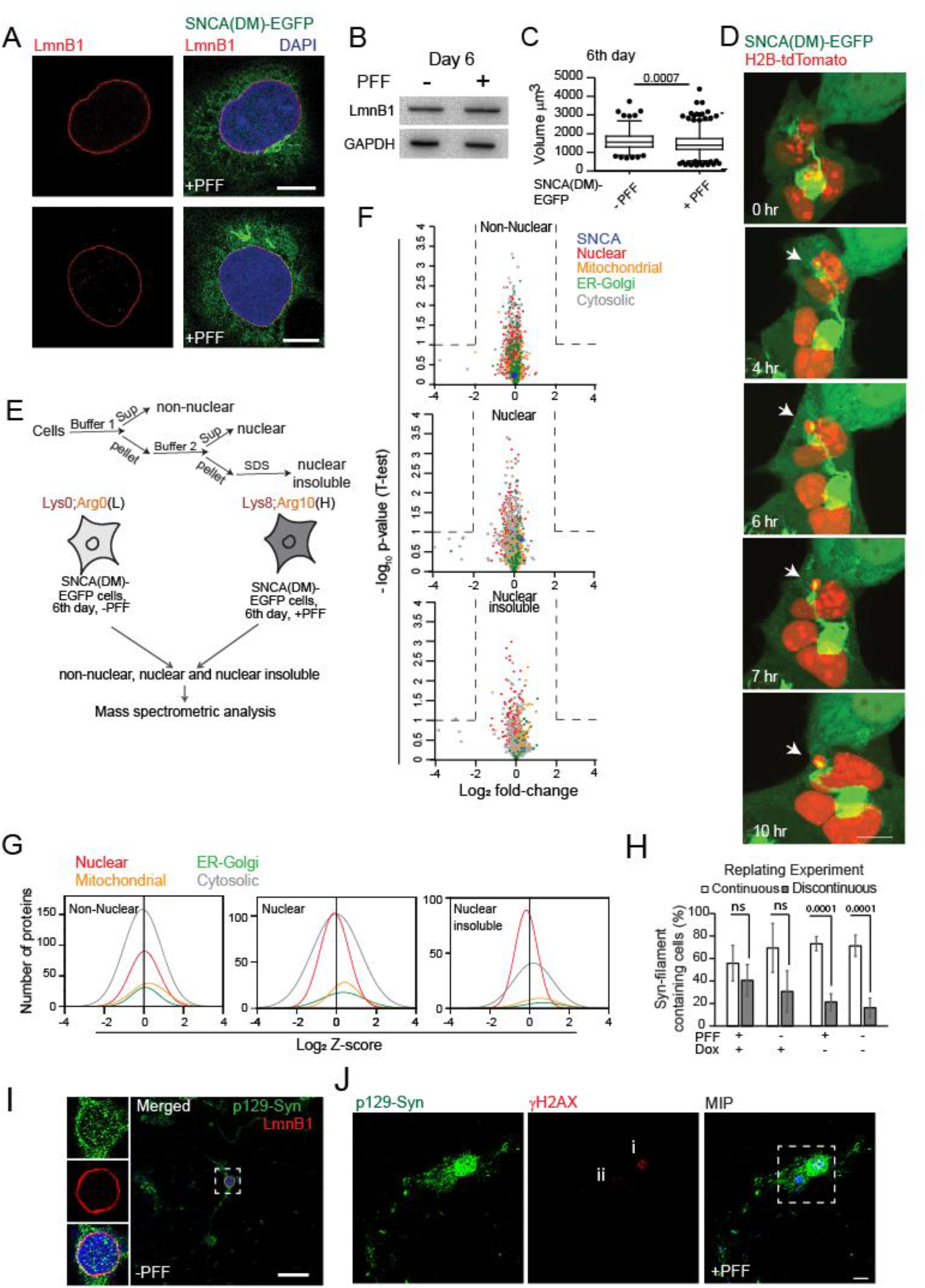
Perinuclear LB-like IBs trigger nuclear lamina deformities. **A**: 6^th^ day PFF treated SNCA(DM)-EGFP cells immunostained for LmnB1. **B:** Western blot analysis of 6^th^ day total fractions of SNCA(DM)-EGFP (-/+) PFF cells with LmnB1 antibody. GAPDH as loading control. **C:** Nuclear volume as estimated by Imaris 8.4 (Bitplane) in 6^th^ day SNCA(DM)-EGFP cells. Mann-Whitney U test. **D:** Live cell microscopy images showing nuclear deformities. SNCA(DM)-EGFP cells transfected with H2B-tdTomato on 3rd day of PFF incubation. Arrows indicate deformity site. **E:** Experimental strategy for Non-Nuclear, Nuclear soluble and Nuclear insoluble fraction preparation. **F:** Volcano plot of 6^th^ day PFF treated SNCA(DM)-EGFP cells. The −log10 (p-value, One-sample *t*-test) plotted against log2 (fold change: SNCA(DM)-EGFP; 6^th^ day +PFF/-PFF) of Non-Nuclear, Nuclear and Nuclear pellet fraction. **G:** Z-score (SNCA(DM)-EGFP, 6^th^ day; Ratio: +PFF/-PFF) distribution of proteins identified by mass spectrometry according to their subcellular localization (as curated by UniProt) in Non-Nuclear, Nuclear and Nuclear insoluble fractions. **H:** Replating experiment as per **Fig. 3F**. Continuous or discontinuous lamina as per LmnB1 staining estimated in fragmented or filamentous IB containing cells and plotted. Error bars – SDs. N=3. Student’s *t*-test. **I:** Primary neurons at 15 DIV were fixed and immunostained with p129-Syn and LmnB1. Insets: zoomed images of the indicated locations. **J:** Maximum intensity Projections (MIP) of larger field for image shown in **Fig. 5J**. Scale bar-10 µm. **Fig. S5A, I-J** – Acetone:Methanol fixation. **Fig. S5D** – Live cell imaging.

**Fig. S6:**
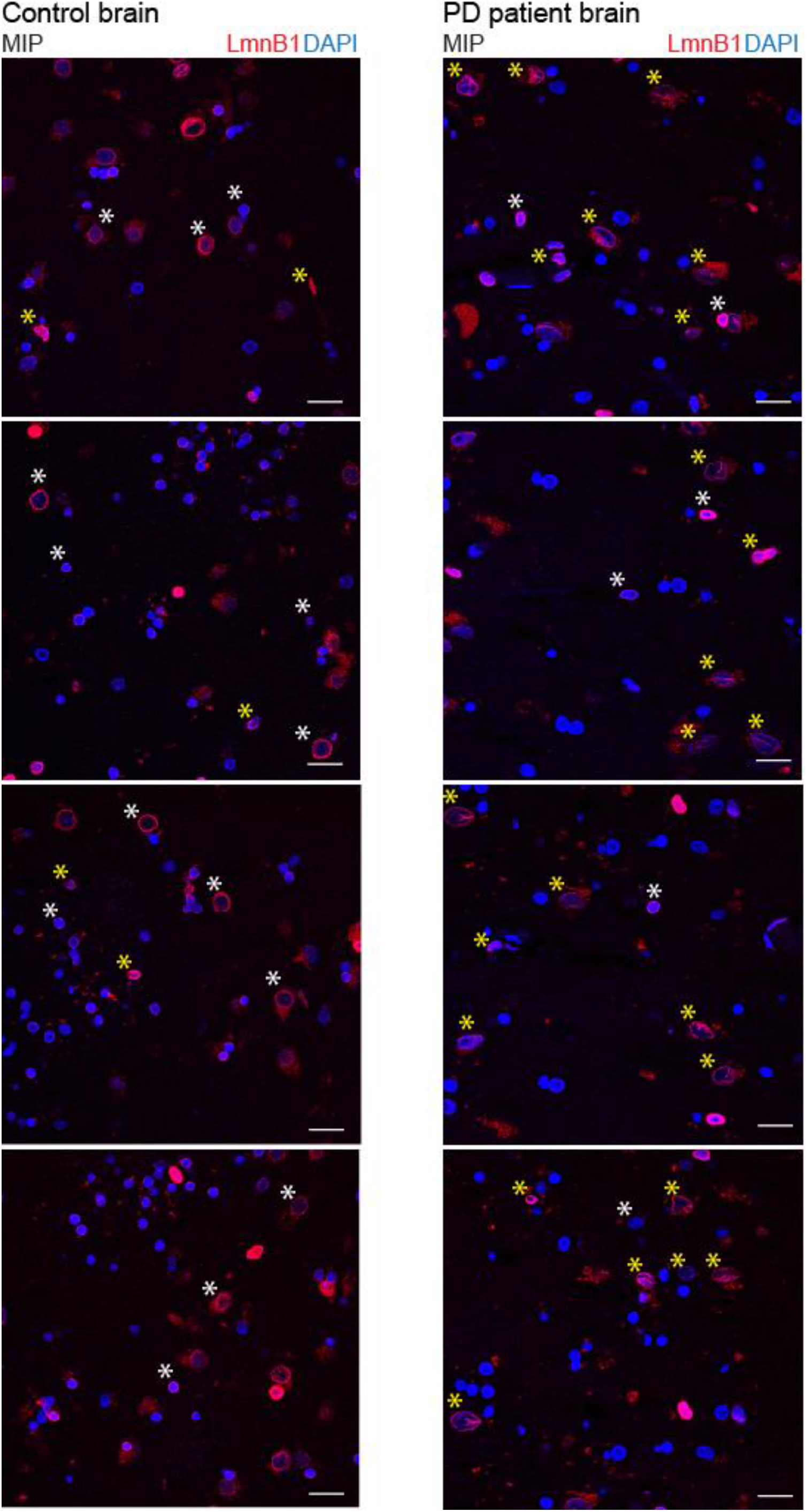
LBs trigger nuclear lamina deformities. MIP of a non-neurological disease (control) and Parkinson’s disease (PD) patients’ brain sections were stained with LmnB1 antibody. insets: zoomed images of box-indicated locations. Scale bar-20 µm. White asterisk – representative nuclei unperturbed lamina; Yellow asterisk – nuclei with lamina disintegrities.

**Fig. S7:**
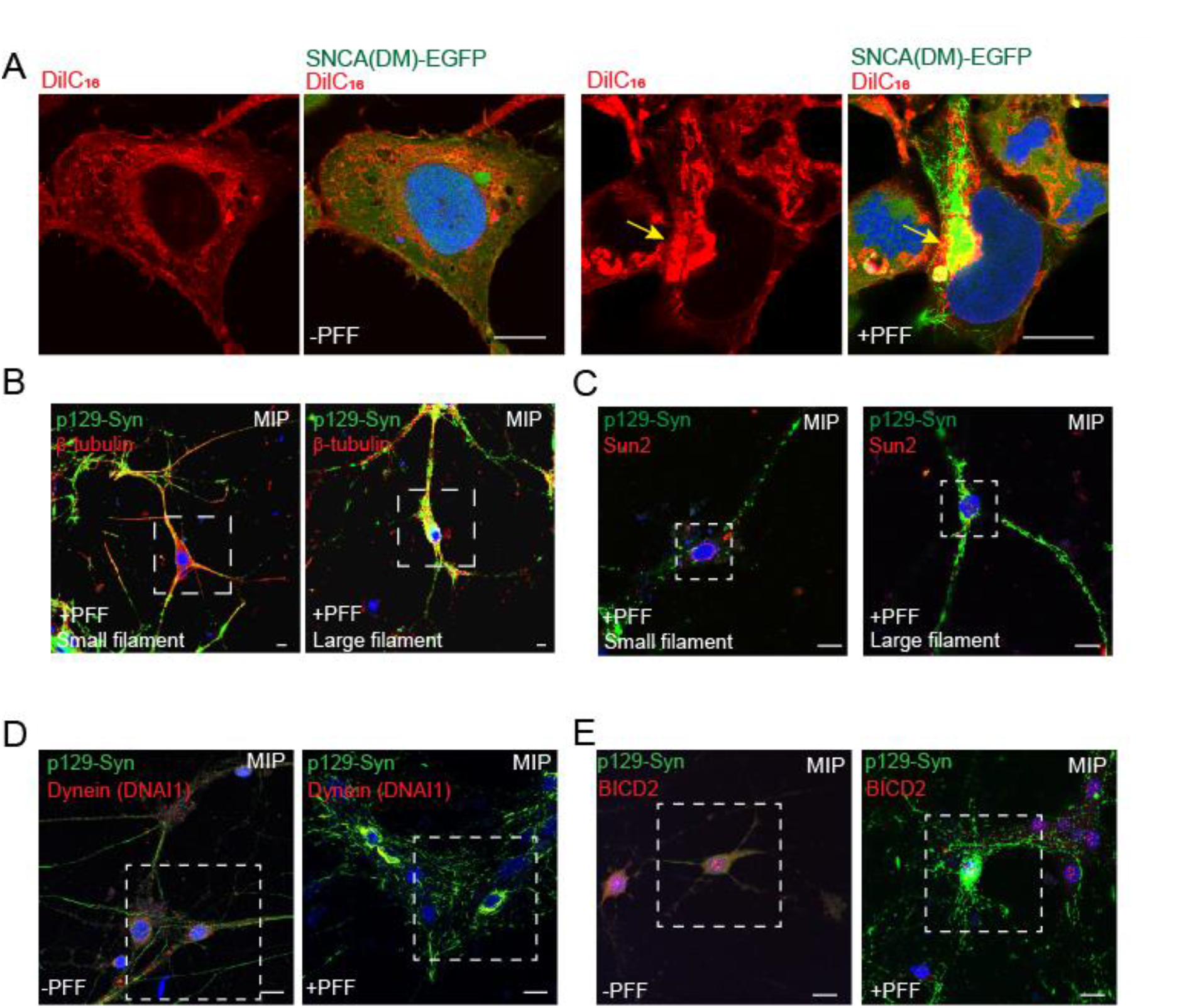
Mis-localization of cytoplasmic motor proteins on LB-like IBs in primary neurons. **A**: Microscopic images of 6^th^ day (-/+) PFF treated SNCA(DM) cells stained with lipid staining dye DiIC16. Yellow arrow indicates Co-localization. **B-E:** MIP of larger field of primary neurons at 15 DIV shown in **Fig. 7B–E**. Dotted box indicates the field of view shown in **Fig. 7B–E**. Scale bar-10 µm. **Fig. S7A** – Paraformaldehyde fixation. **Fig. S7B–E** – Acetone:Methanol fixation.

**Fig. S8:**
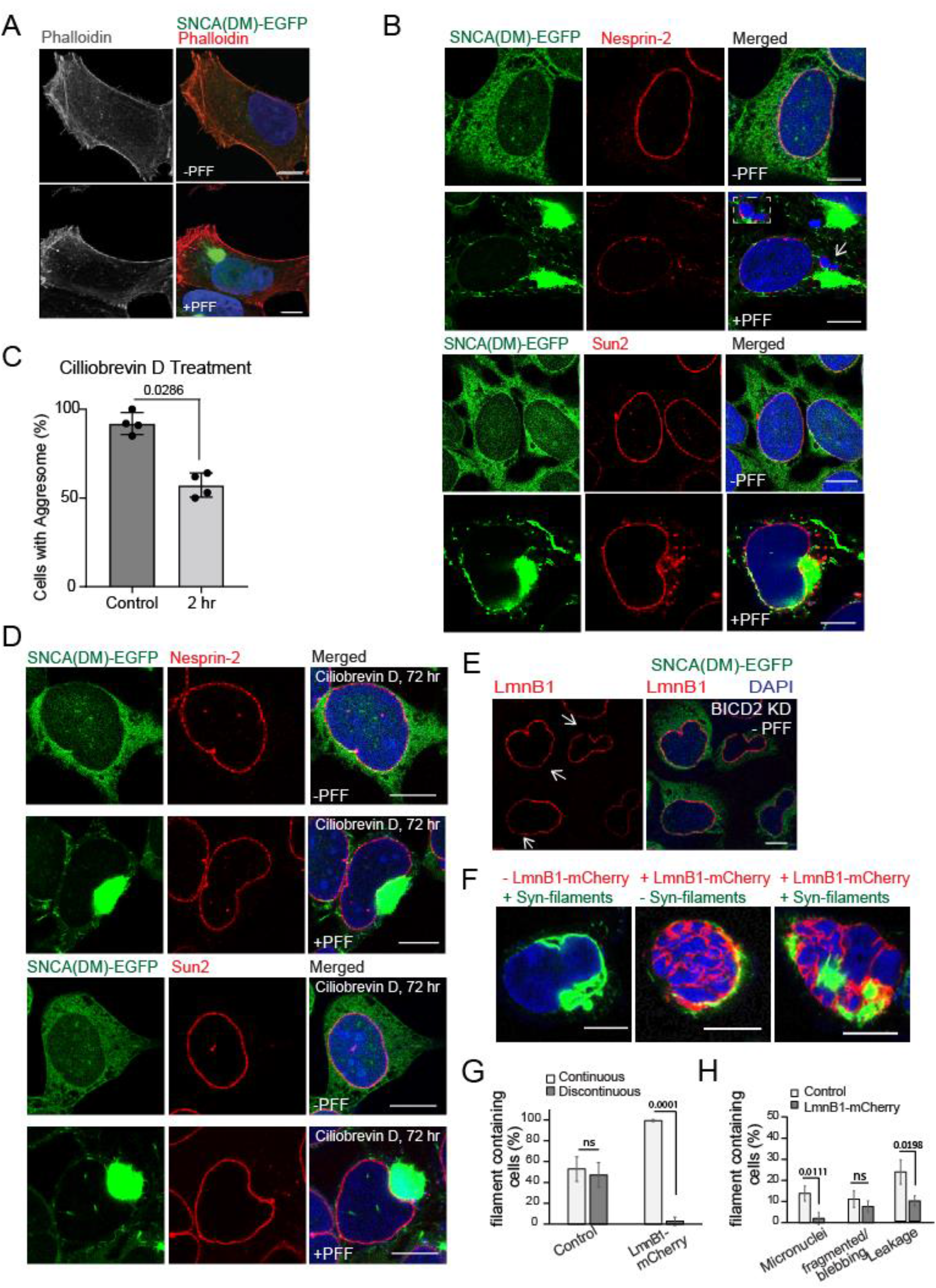
Altered cytoskeleton-forces in LB-like IB containing cells trigger NE-injuries. **A**: Maximum Intensity projection (MIP) of microscopic images of 6^th^ day (-/+) PFF treated SNCA(DM)-EGFP cells stained with Phalloidin-red. **B:** Microscopic images shown in **Fig. 8B** – Images of fluorescent proteins/antibodies excited by different wavelengths of lasers are shown separately. **C:** No. of aggresome containing cells counted in control and Ciliobrevin D treated SNCA(DM)-EGFP cells and plotted. Student’s *t*-test (N=3). **D:** Microscopic images shown in **Fig. 8G** – Images of fluorescent proteins/antibodies excited by different wavelengths of lasers are shown separately. **E:** 6th day without PFF treatment, SNCA (DM)-EGFP cells immunostained with LmnB1 after 72 hr BICD2 knockdown (KD). White arrows indicate lamina gaps. **F:** PFF treated SNCA(DM)-EGFP cells transfected with LmnB1-mCherry on 5^th^ day and imaged on 6^th^ day. **G-H:** Deformities in nuclear morphology and appendages as per LmnB1-mCherry and DAPI staining plotted. Approx. 50 cells counted in each experiment. N=3, Student’s *t*-test. Scale bar – 10 µm. **Fig. S7A** – Paraformaldehyde fixation. **Fig. S7B, D-F** – Acetone:Methanol fixation.

**Fig. S9:**
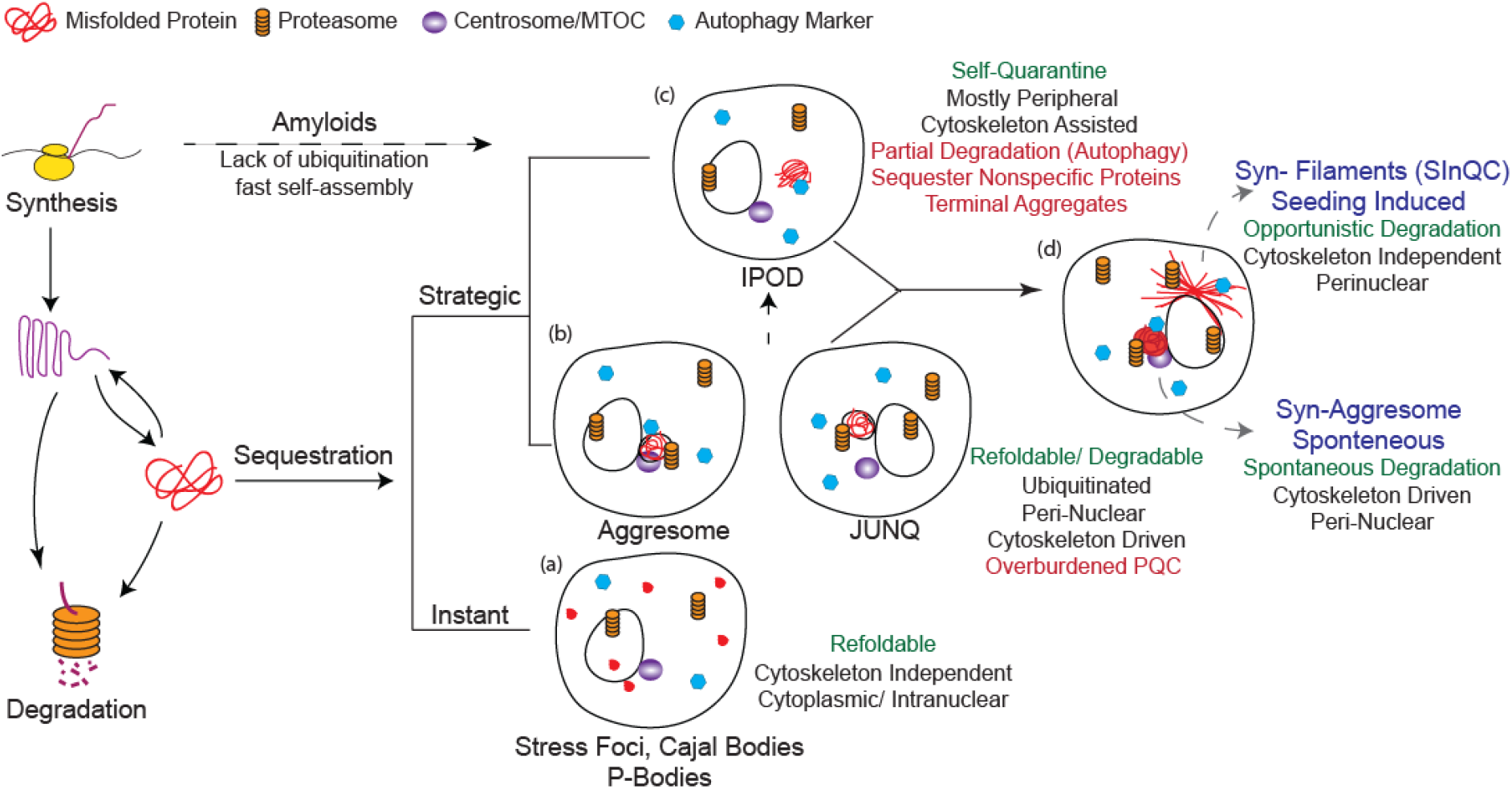
Different strategies to sequester misfolded protein load in the cell. (**a**) Misfolding-prone proteins are sequestered into multiple reversible stress foci in cytoplasm and nucleus instantaneously after acute stress such as heat shock. (**b**) Upon continuous stress, misfolding prone proteins are ubiquitinated and directed to perinuclear sites in cytoskeleton driven manner. These are called aggresomes or JUNQs depending on their position on MTOC or their proximity to perinuclear endoplasmic reticulum. Both aggresomes and JUNQs serve as sites for refolding or degradation of misfolded proteins depending on their recognition by PQC-factors. **(c**) Upon saturation of the capacity of aggresomes/JUNQs, further increased load of misfolded proteins is sequestered into peripheral sites known as IPODs. Proteins accumulated into IPODs are terminally aggregated and avoid non-specific interactions with cellular proteins. IPODs are occasionally degraded by autophagy machinery or diluted through division in mitotic cells. Amyloids possessing faster self-polymerization properties are proposed to directly accumulate into IPODs bypassing aggresomes/JUNQs. **(d**) Misfolding prone α-Synuclein spontaneously accumulate into **Syn-aggresomes** for continuous turnover and into **Syn-filaments** upon seeding. Syn-filaments deposits into perinuclear LBs upon maturation and remain opportunistically UPS-degradable via aggresomes. Perinuclear LBs also share the self-sequestration properties of IPODs, remain less interactive and seclude misfolding prone α-Synuclein from the rest of the proteome. Thus, perinuclear LBs represent a unique homeostatic inclusion body sharing properties of both aggresomes and IPODs and we term these as **SInQCs – Seeding based Inclusions for Quality Control**.

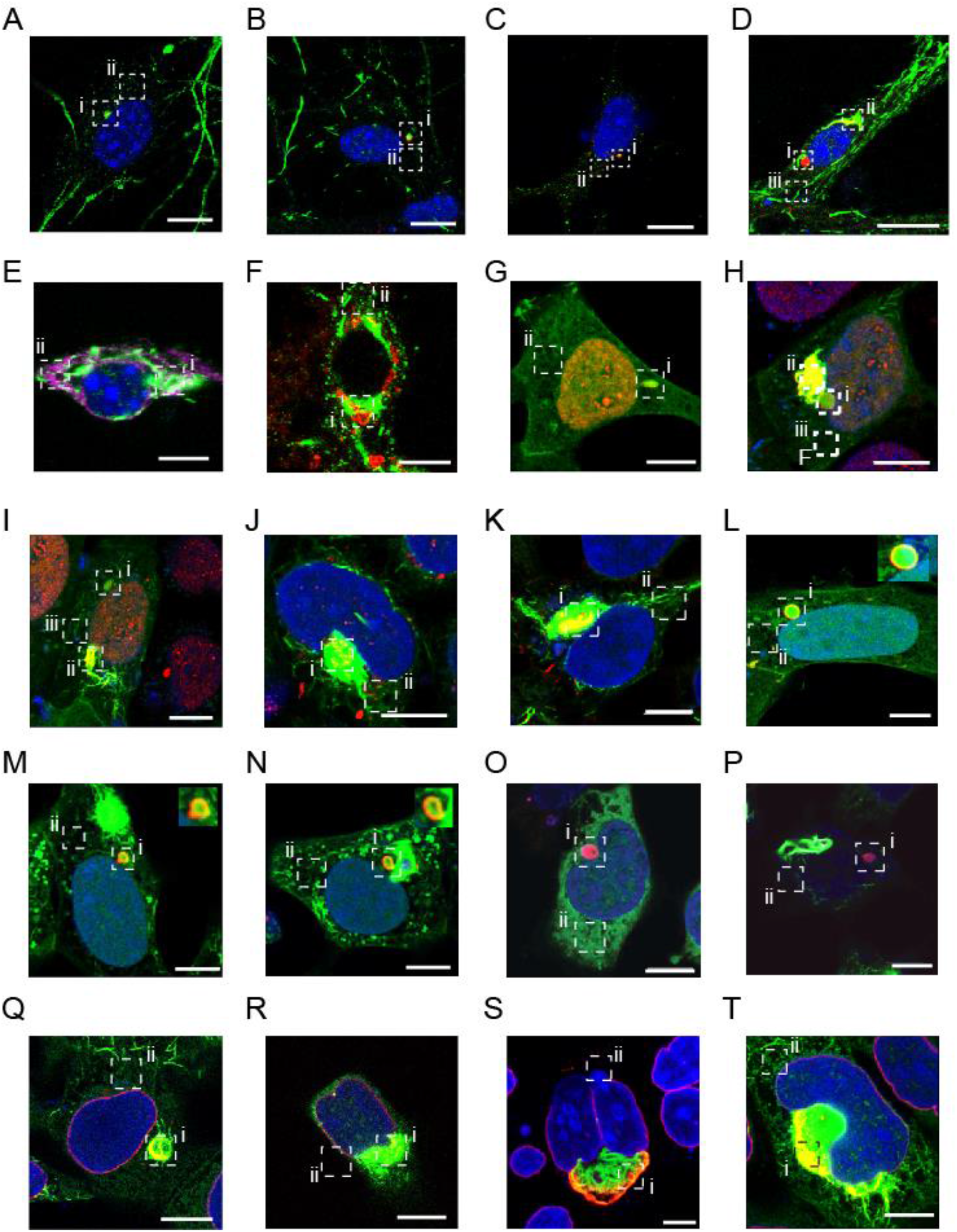
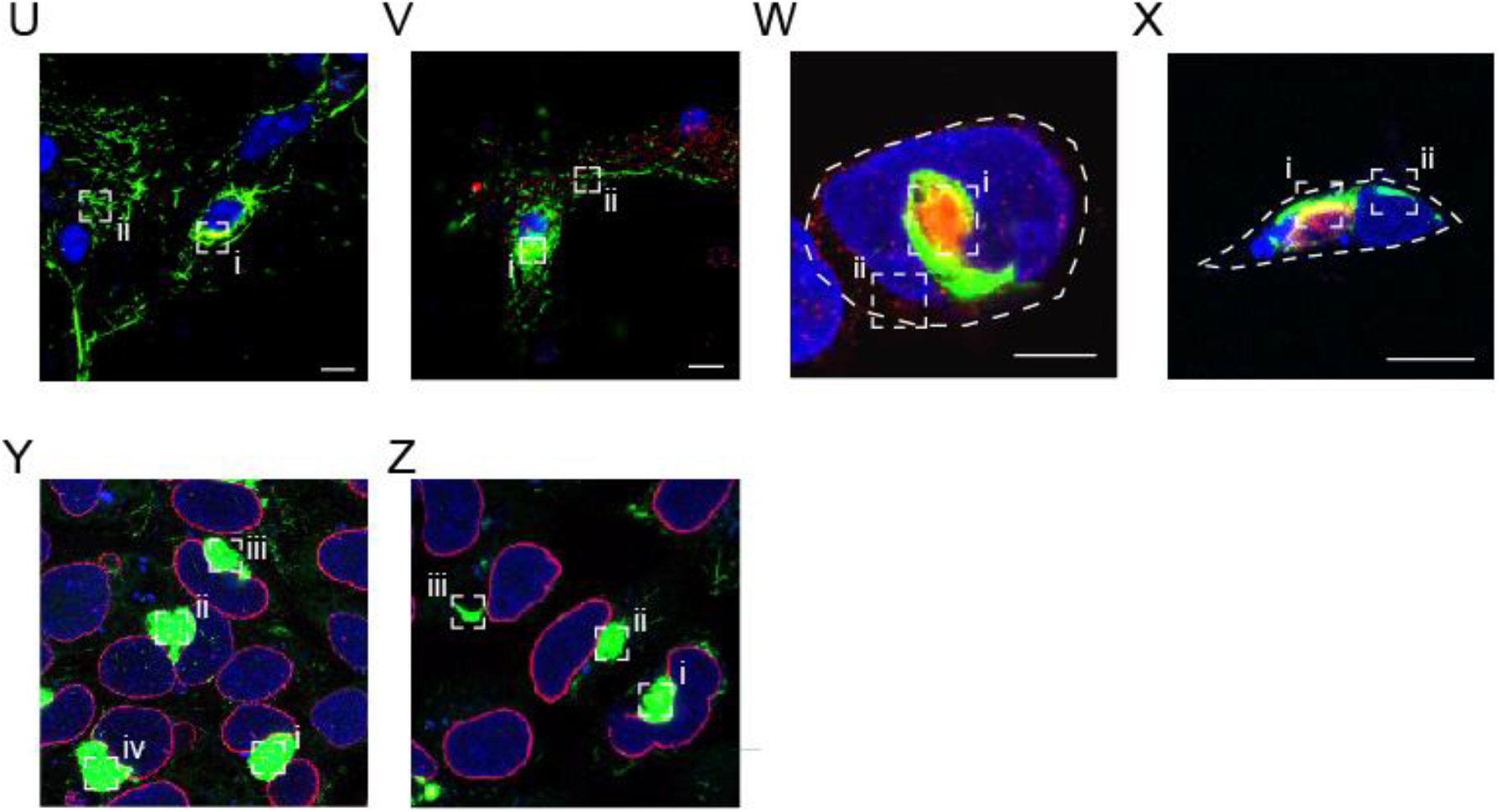
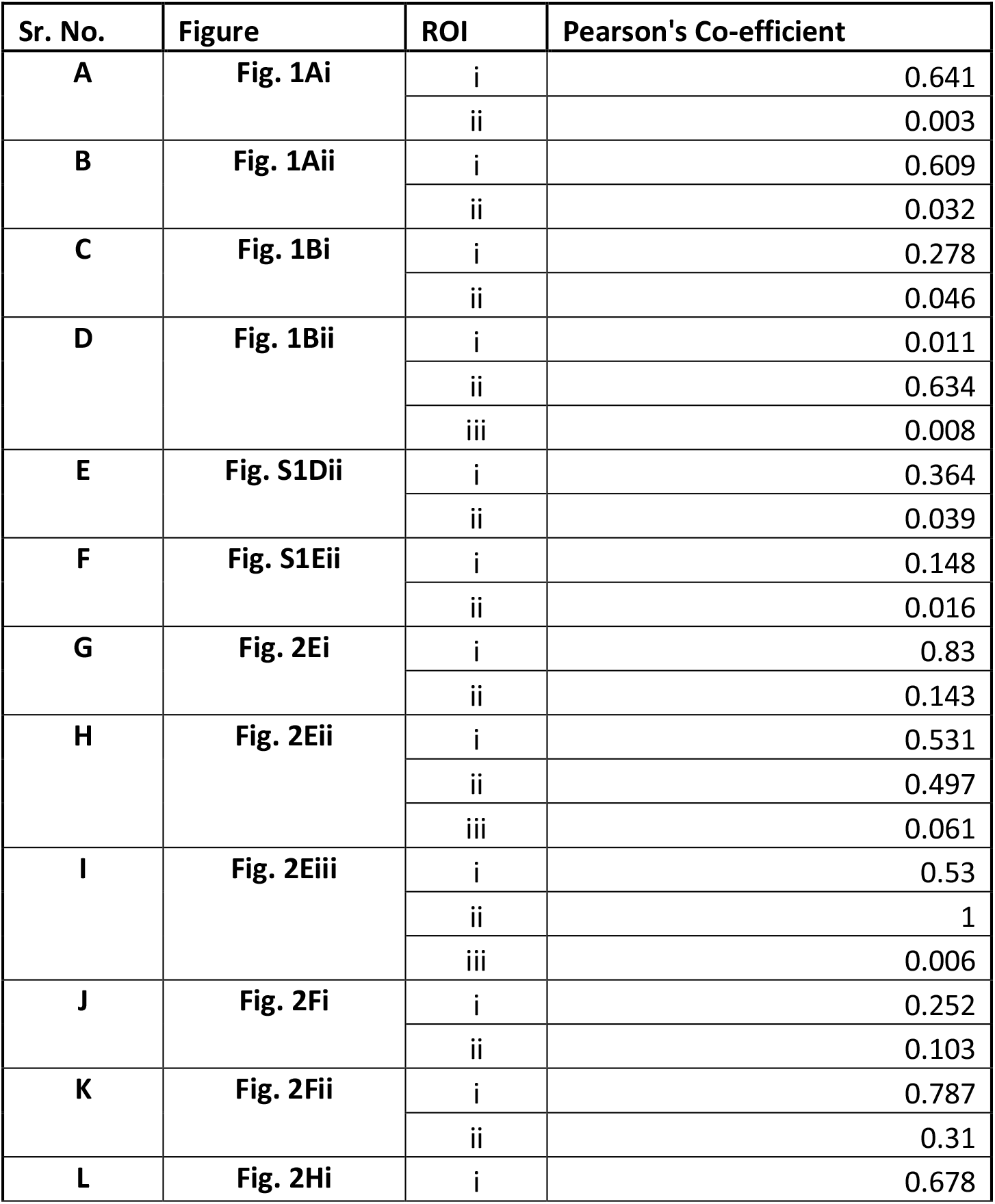
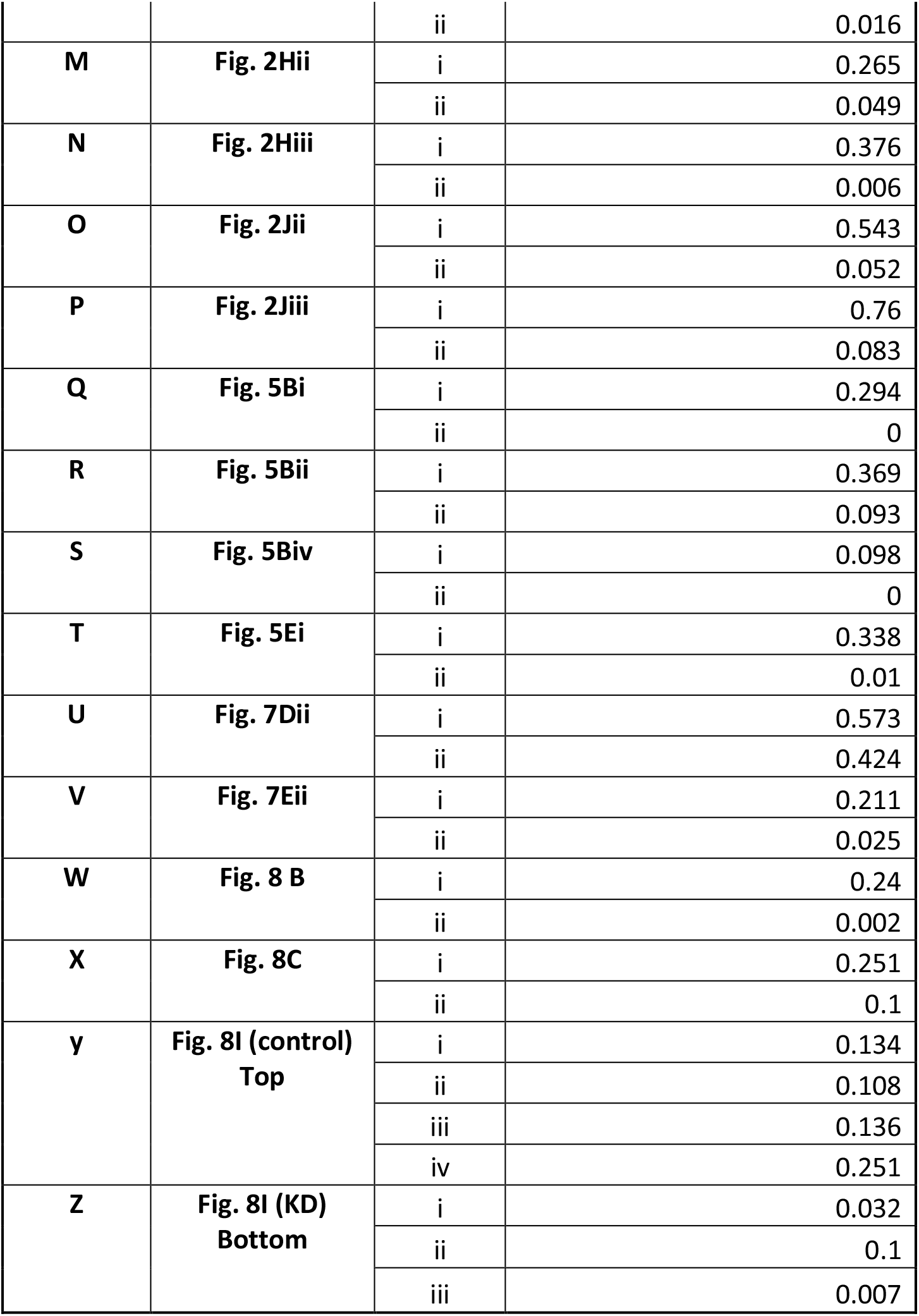
Supplementary note: Colocalization figures and table. Pearson’s co-localization coefficient was calculated for the indicated ROI by JACoP algorithm in ImageJ (91).

**Table S1. Z-scores distribution for total, soluble and insoluble fractions**.

Z-scores distribution of fold change of total, soluble and insoluble fractions.

**Table S2. Fold change distribution of PQC-proteins**.

Log2 (fold change: SNCA(DM)-EGFP, 6day +PFF/-PFF) distribution of PQC-proteins on 6th day.

**Table S3. One sample t-test calculation of total, soluble and insoluble fractions**.

P-values calculated by one sample t-test of fold change (HEK293T SNCA-DM-EGFP, + PFF/-PFF) for Total fraction at day 6.

**Table S4. Fold change distribution of heat shock proteins in primary neurons**.

Log2 (fold change: SNCA(DM)-EGFP-NLS, 6day, +PFF (Heat Shock/ Non-Heat Shock) and SNCA(DM)-EGFP-NLS, 6day, -PFF (Heat Shock/ Non-Heat Shock) distribution of PQC-proteins on 6th day

**Table S5. Z-scores distribution for NNF, NS and NI fraction**.

Z-scores distribution of fold change (HEK293T SNCA-DM-EGFP, + PFF/-PFF) at Day 6 of nuclear insoluble fraction.

**Table S6. One sample t-test calculation of NNF, NS and NI fraction**.

P-values calculated by one sample t-test of fold change (HEK293T SNCA-DM-EGFP, + PFF/-PFF) at day 6 for non-nuclear fraction.

**Movie S1. Syn-filaments at different stages of biogenesis**.

**Movie S2. Syn-filament containing cell undergoing mitotic cell division**.

**Movie S3. Detachment of Syn-filament containing cell from plate surface**.

**Movie S4. Nuclear hernia-like blobs were captured in Syn-filament containing cell**.

**Movie S5. Nuclear rupture in Syn-filament containing cells**.

